# Shadow enhancers suppress input transcription factor noise through distinct regulatory logic

**DOI:** 10.1101/778092

**Authors:** Rachel Waymack, Alvaro Fletcher, German Enciso, Zeba Wunderlich

## Abstract

Shadow enhancers, groups of seemingly redundant enhancers, are found in a wide range of organisms and are critical for robust developmental patterning. However, their mechanism of action is unknown. We hypothesized that shadow enhancers drive consistent expression levels by buffering upstream noise through a separation of transcription factor (TF) inputs at the individual enhancers. By measuring transcriptional dynamics of several *Kruppel* shadow enhancer configurations in live *Drosophila* embryos, we showed individual member enhancers act largely independently. We found that TF fluctuations are an appreciable source of noise that the shadow enhancer pair can better buffer than duplicated enhancers. The shadow enhancer pair is uniquely able to maintain low levels of expression noise across a wide range of temperatures. A stochastic model demonstrated the separation of TF inputs is sufficient to explain these findings. Our results suggest the widespread use of shadow enhancers is partially due to their noise suppressing ability.

## Introduction

The first evidence that transcription occurred in bursts, as opposed to as a smooth, continuous process, was observed in *Drosophila* embryos, where electron micrographs showed that even highly transcribed genes had regions of chromatin lacking associated transcripts between regions of densely associated nascent transcripts (Miller & McKnight, 1979). As visualization techniques have improved, it is increasingly clear that transcriptional bursting is the predominant mode of expression across organisms from bacteria to mammals (Dar, et al.,2012; Sanchez & Golding, 2013; Zenklusen, et al., 2008; Fukaya, et al., 2016). These bursts of transcriptional activity, separated by periods of relative silence, have important implications for cellular function, as mRNA numbers and fluctuations largely dictate these quantities at the protein level (Csardi, et al., 2015; Hansen, et al., 2018). Such fluctuations in regulatory proteins, like TFs and signaling molecules, can propagate down a gene regulatory network, significantly altering the expression levels or noise of downstream target genes (Blake, et al., 2003).

Given that protein levels fluctuate and that these fluctuations can cascade down regulatory networks, this raises the question of how organisms establish and maintain the precise levels of gene expression seen during development, where expression patterns can be reproducible down to half-nuclear distances in *Drosophila* embryos (Dubuis, et al., 2013; Gregor, et al., 2007). Many mechanisms that buffer against the noise inherent in gene expression or stemming from genetic or environmental variation have been observed (Lagha, et al., 2012; Stapel, et al., 2017; Raj et al., 2010). For example, organisms use temporal and spatial averaging mechanisms and redundancy in genetic circuits to achieve the precision required for proper development (Stapel, et al., 2017; Raj, et al., 2010; Erdman, et al., 2009; Lagha, et al., 2012).

Here, we propose that shadow enhancers may be another mechanism by which developmental systems manage noise (Barolo, S., 2012). Shadow enhancers are groups of two or more enhancers that control the same target gene and drive overlapping spatiotemporal expression patterns (Barolo, S., 2012). Shadow enhancers are found in many organisms, from insects to plants to mammals, and are strongly associated with developmental genes (Cannavo, et al., 2016; Osterwalder, et al., 2018; Garnett, et al., 2012; Bomblies, et al., 1999). These seemingly redundant enhancers have been shown to be critical for proper gene expression in the face of both environmental and genetic perturbations, which may exacerbate fluctuations in upstream regulators (Frankel, et al., 2010; Osterwalder, et al. 2018; Perry, et al., 2010; Cheung & Ma, 2015, Chen, et al., 2015). However, shadow enhancers’ precise mechanism of action is still unknown. Others have proposed that having multiple enhancers controlling the same promoter ensures a critical threshold of gene expression is reached, perhaps by reducing the effective “failure rate” of the promoter (Lam, et al., 2015; Perry, et al., 2011). An alternative, but not mutually exclusive, possibility is that shadow enhancers ensure precise expression by buffering noise in upstream regulators. Several studies suggest that individual enhancers of a shadow enhancer group tend to be controlled by different sets of TFs, which we call a “separation of inputs” (Wunderlich, et al., 2015; Cannavo, et al., 2016; Ghiasvand, et al., 2011). We hypothesize that this separation allows shadow enhancers to buffer against fluctuations in TF levels.

The *Drosophila* gap gene *Kruppel* (*Kr*) provides a useful system in which to address the mechanisms of action of shadow enhancers. During early embryogenesis, *Kr* is controlled by the activity of two enhancers, proximal and distal, that are each activated by different sets of TFs (Figure 1A; Wunderlich, et al., 2015). Here we focus on differences in activation, as the key repressors of *Kr*, *knirps* and *giant*, are likely to regulate both enhancers. *Kr* expression during this time is critical for thorax formation, and like the other gap genes in the *Drosophila* embryo, has quite low noise (Preiss, et al., 1985; Dubuis, et al., 2013). By measuring live mRNA dynamics, we can use the *Kr* system in *Drosophila* embryos to assess whether and how shadow enhancers act to buffer noise and identify the sources of noise in the developing embryo.

**Figure 1:**
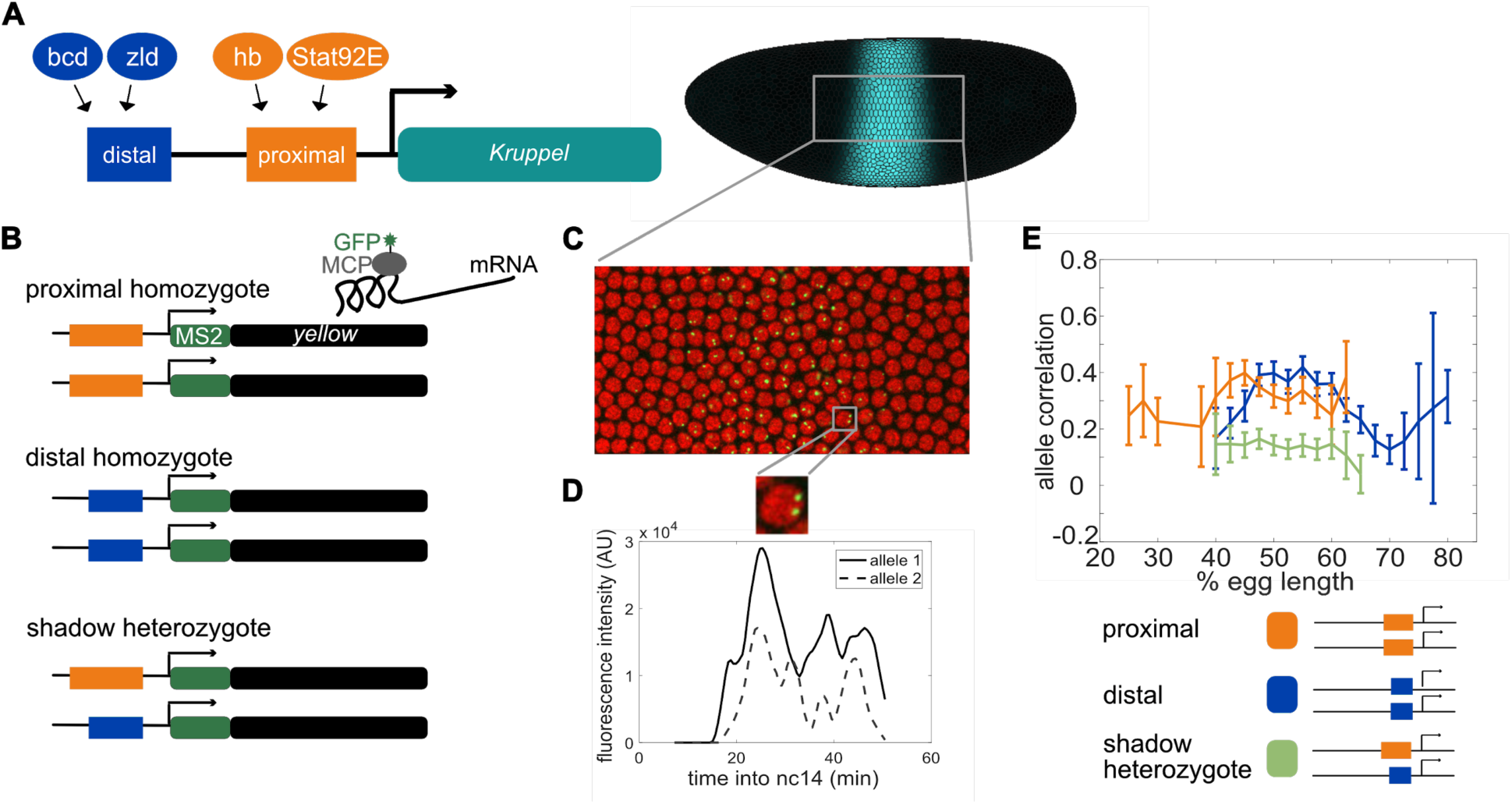
Dual allele imaging shows the individual *Kruppel* enhancers drive largely independent transcriptional dynamics. **A**. Schematic of the endogenous *Kruppel* locus with distal (blue) and proximal (orange) shadow enhancers driving *Kr* (teal) expression in the central region of the embryo. Known transcriptional activators of the two enhancers are shown. **B**. Schematics of single enhancer reporter constructs driving expression of MS2 and a *yellow* reporter. When transcribed, the MS2 sequence forms stem loops that are bound by GFP-tagged MCP expressed in the embryos. Proximal embryos have expression on each allele controlled by the 1.5kb proximal enhancer at its endogenous spacing from the *Kr* promoter, while distal embryos have expression on each allele controlled by the 1.1kb distal enhancer at the same spacing from the *Kr* promoter. Shadow heterozygote embryos have expression on one allele controlled by the proximal enhancer and expression on the other allele controlled by the distal enhancer. **C**. Still frame from live imaging experiment where nuclei are red circles and active sites of transcription are green spots. MCP-GFP is visible as spots above background at sites of nascent transcription (Garcia, et al., 2013). **D.** The fluorescence of each allele in individual nuclei can be tracked across time as a measure of transcriptional activity. Graph shows a representative trace of transcriptional activity of the two alleles in a single nucleus across the time of nc14. These traces are used to calculate the correlation of allele activity in each nucleus. Correlation values are grouped by position of the nucleus along the egg length and averaged across all imaged nuclei in all embryos of each construct. **E**. Graph of average correlation between the two alleles in each nucleus as a function of egg length. 0% egg length corresponds to the anterior end. Error bars indicate 95% confidence intervals. The shadow heterozygotes have much lower allele correlation than either homozygote, demonstrating that the individual shadow enhancers drive nearly independent transcriptional activity and that upstream fluctuations in regulators are a significant driver of transcriptional bursts. The total number of nuclei used in calculations for each construct by AP bin are given in Supplementary Table 2.

To test our hypothesis, we measured live mRNA dynamics driven by either single *Kr* enhancer, duplicated enhancers, or the shadow enhancer pair and compared the dynamics and noise associated with each. We showed that the individual *Kr* enhancers can act largely independently in the same nucleus while identical enhancers display correlated activity. We constructed a simple mathematical model to describe this system and found that TF fluctuations are necessary to reproduce the correlated activity of identical enhancers in the same nucleus. Using this model, we also found that the lower expression noise driven by the shadow enhancer pair compared to either duplicated enhancer is a natural consequence of the separation of TF inputs. Experimentally, we found the shadow enhancer pair achieves lower noise through decreases in both intrinsic and extrinsic sources of noise. Additionally, the shadow enhancer pair is uniquely able to maintain low levels of expression noise across a wide range of temperatures. We suggest that this noise suppression ability is one of the key features that explains the prevalence of shadow enhancers in developmental systems.

## Results

### Individual enhancers in the shadow enhancer pair act nearly independently within a nucleus

To test our hypothesis that the separation of inputs between *Kruppel’s (Kr)* shadow enhancers provides them with noise-buffering capabilities, we needed to first test the ability of each enhancer to act independently. If variability in gene expression is driven primarily by fluctuations in upstream factors, the shadow enhancer pair, whose individual enhancers are controlled by different sets of TFs, could provide a form of noise buffering. Conversely, variability in upstream regulators may be low enough in the developing embryo that these fluctuations are not the primary driver of downstream expression noise. If this were the case, the separation of inputs is unlikely to be a key requirement of shadow enhancer function.

To investigate these possibilities, we measured and compared the correlation of allele activity in homozygous or heterozygous embryos that carry two reporter genes. *Proximal homozygotes* contained the proximal enhancer driving a reporter, inserted in the same location on both homologous chromosomes, and *distal homozygotes* similarly had the distal enhancer driving reporter expression on both homologous chromosomes (Figure 1B). We also made heterozygous embryos, called *shadow heterozygotes*, which had one proximal and one distal reporter, again in the same location on both homologous chromosomes. To measure live mRNA dynamics and correlations in allele activity, we used the MS2-MCP reporter system (Figure 1C, D). This system allows the visualization of mRNAs that contain the MS2 RNA sequence, which is bound by an MCP-GFP fusion protein (Bertrand, et al., 1998). In the developing embryo, only the site of nascent transcription is visible, as single transcripts are too dim, allowing us to measure the rate of transcription (Garcia, et al., 2013; Lucas, et al., 2013). In blastoderm-stage embryos with two MS2 reporter genes, we can observe two distinct foci of fluorescence corresponding to the two alleles (Figure 1D), in line with previous results that suggest there are low levels of transvection at this stage (Lim, et al., 2018; Fukaya & Levine, 2017). To confirm our ability to distinguish the two alleles, we imaged transcription in embryos hemizygous for our reporter constructs, which only show one spot of fluorescence per nucleus. Our counts of active transcription sites in homozygous embryos correspond well to the expected value calculated from hemizygous embryos (Supplemental Figure 1). Therefore, we are able to measure the correlation of allele activity, though we cannot identify which spot corresponds to which reporter.

We predicted that if variability in gene expression is driven by fluctuations in input TFs, we would observe lower correlations of allele activity in shadow heterozygotes than in either the proximal or distal homozygotes. However, if global factors affecting both enhancers dominate, there would be no difference in allele activity correlations. During the ∼1 hour of nuclear cycle 14 (nc14) we found that allele activity is more than twice as correlated in both proximal and distal homozygotes than in shadow heterozygote embryos at 47-57% egg length, which encompasses the central region of *Kr* expression during this time period (Figure 1). This indicates not only that the individual member enhancers of the shadow enhancer pair can act largely independently in the same nucleus, but that differential TF inputs are the primary determinants of transcriptional bursts in this system. Notably, heterozygotes still show marginal allele correlation, indicating that some correlation is induced by either shared input TFs or factors that affect transcription globally. The independence of individual *Kr* enhancers allows for the possibility that shadow enhancers can act to buffer noise by providing separate inputs to the same gene expression output.

### Transcription factor fluctuations are required for the observed differences in the correlations of enhancer activity

To explore the conditions needed for the two *Kr* enhancers to act nearly independently within the same nucleus, we generated a simple model of enhancer-driven dynamics. We considered an enhancer E that interacts with a transcription factor T, which together bind to the promoter to form the active promoter-enhancer complex C (Figure 2A). When the promoter is bound by the enhancer, it drives the production of mRNA. Since the MS2 system only allows us to observe mRNA at the site of transcription, we modeled the diffusion of mRNA away from the transcription site as decay. The transcription factor T is produced in bursts of *n* molecules at a time, and it degrades linearly. For simplicity, the transcription factor T is an abstraction of the multiple activating TFs that interact with the enhancer, and T corresponds to a different set of TFs for the proximal and distal enhancer. This nonlinear model generalizes the linear model by Bothma et al. (Bothma et al., 2015) by explicitly taking into account the presence of TFs.

**Figure 2:**
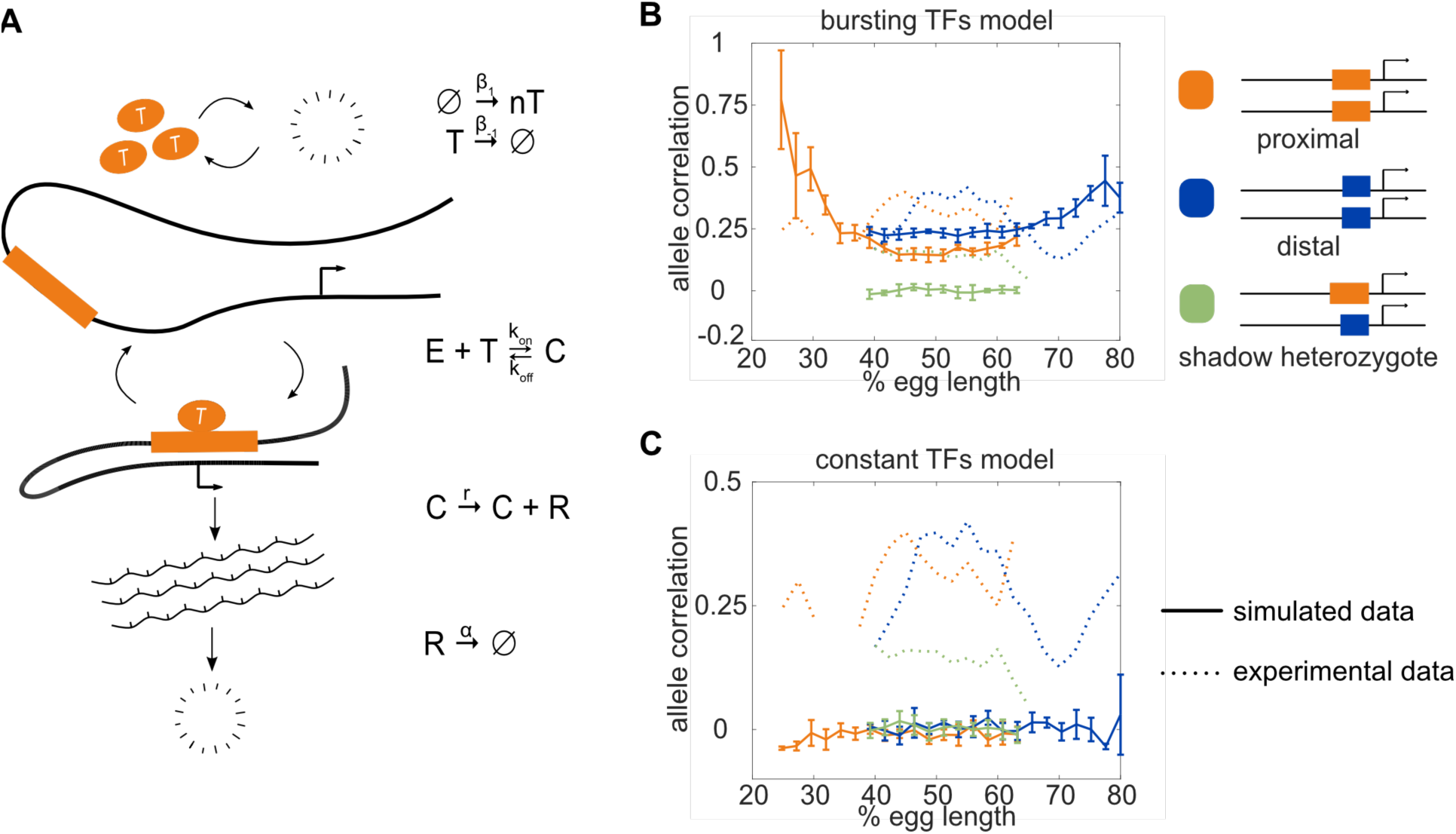
Model of enhancer-driven dynamics demonstrates TF fluctuations are required for correlated reporter activity. To investigate the factors required for the observed correlated behavior of identical enhancers and largely independent behavior of the individual enhancers, we developed a simple stochastic model of enhancer-driven transcription. **A**. Schematic of model of transcription driven by a single enhancer (the *bursting TFs* model). For each enhancer, we assume there is a single activating TF, T, that appears in bursts of size *n* molecules at a rate β_1_, which varies by the position in the embryo. TFs degrade linearly at rate β_-1_. When present, T can bind the enhancer, E, to form a transcriptionally active complex, C, at a rate *k_on_* and dissociates at rate *k_off_*. This complex then produces mRNA at an experimentally determined rate *r* that degrades at an experimentally determined rate, α. **B**. The bursting TFs model is able to recapitulate the experimentally observed pattern of allele correlation. We plot the correlation between the two alleles in a nucleus as a function of egg length. Simulated data is created using the lowest energy parameter set for each enhancer. The data shown is the average of five simulated embryos that have 80 transcriptional spots per AP bin. In B and C simulated data are shown by solid lines, experimental data are shown by dotted lines. **C**. The constant TFs model fails to recapitulate the experimentally observed pattern of allele correlation. Without TF fluctuations, both heterozygous and homozygous embryos display independent allele activity. Error bars in B and C represent 95% confidence intervals.

We estimated some model parameters directly from experimental data and others by fitting using simulated annealing. The mRNA degradation parameter *α* and production parameter *r* were measured directly from fluorescence data without any input from the model (see Methods for details). The remaining parameters were first estimated using mathematical analysis, then fine-tuned using simulated annealing. We found separate parameter sets for the proximal and distal enhancers that, when used to simulate transcription, fit the experimentally measured characteristics of the transcriptional traces, including transcription burst size, frequency, and duration, as well as the total mRNA produced (Supplementary Figure 2).

We hypothesized that a model that lacks fluctuations in the input TFs could not recapitulate the high correlation of transcriptional activity in homozygotes versus the low correlation in heterozygotes. To test this hypothesis, we generated another model of TF production. We call our original model described above *bursting TFs*. The other model is one in which TF numbers are constant over time, which we call *constant TFs* and is equivalent to the model in (Bothma et al., 2015). If the difference in transcription correlation between homozygotes and heterozygotes is due to fluctuating numbers of TFs, we expected that the bursting TFs model will recapitulate this behavior, while the constant TFs model will not. However, if the constant TFs model is also able to recapitulate the observed difference in correlations, then the correlations are likely a consequence of the identical enhancers simply being regulated by the same set of TFs.

For each model, we used the 10 best parameter sets to simulate transcriptional activity in homozygote and heterozygote embryos and analyzed the resulting allele correlations. We found that the bursting TFs model always produced results in which both homozygote allele correlations are significantly higher than the heterozygote, as observed experimentally (Figure 2B). None of the best fitting parameter sets for the constant TF model were able to produce the experimentally-observed behavior and always resulted in similar correlations for the homozygote and heterozygote embryos (Figure 2C). Therefore, in our minimalist model of enhancer-driven transcription, the presence of TF fluctuations is required for the observed differences in allele correlation. These results also demonstrate the advantage of using a single generic TF for each enhancer. By abstracting away TF interactions, we reduced the complexity and number of parameters in the model, which allowed us to explore the relationship between TF production and allele correlation.

### The shadow enhancer pair drives less noisy expression than enhancer duplications

Since the individual *Kr* enhancers can act independently, we wanted to further test whether this separation of inputs enables the shadow enhancer pair to provide more stable gene expression output. We compared the noise in expression driven by the shadow enhancer pair to that driven by two copies of either the distal or proximal enhancer (Figure 3). If the shadow enhancer pair drives lower noise, this observation, combined with our finding of enhancer independence, strongly suggests that the shadow enhancer pair reduces variability and mediates robustness by buffering fluctuations in upstream regulators. Alternatively, if duplicated enhancers drive similar levels of expression noise, this suggests that a separation of inputs is not critical for shadow enhancer’s function and that shadow enhancers mediate robustness through a different mechanism, such as ensuring a critical threshold of expression is met (Lam, et al., 2015; Perry, et al., 2011).

**Figure 3:**
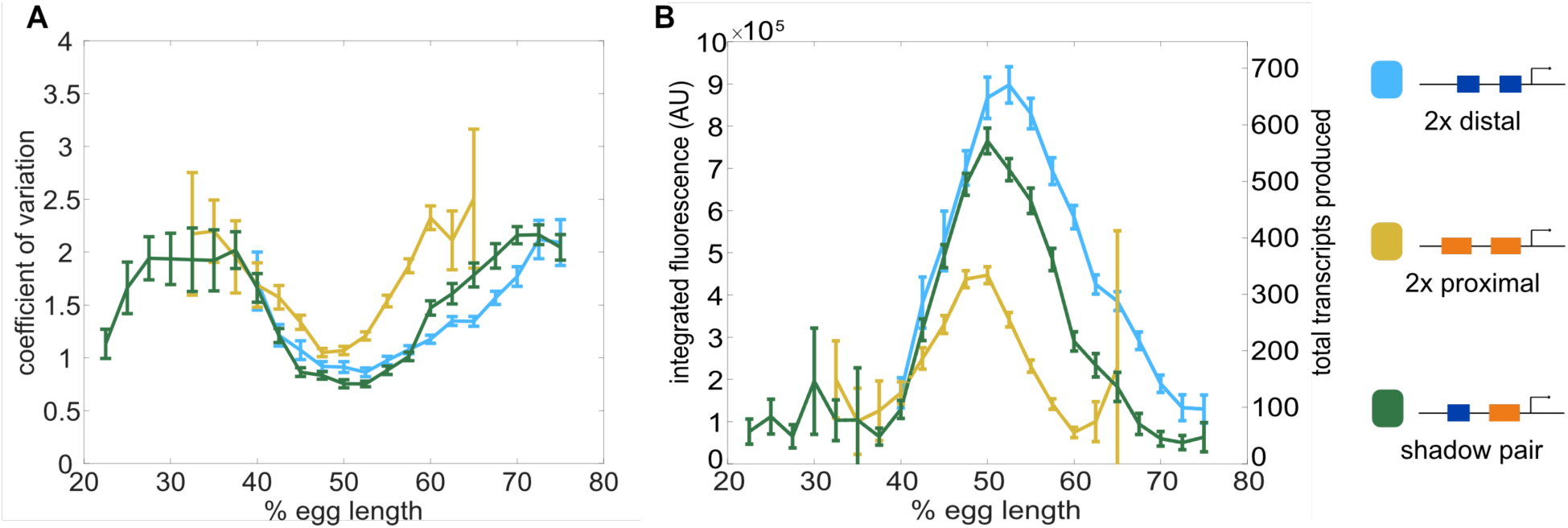
Shadow enhancer pair produces lower expression noise than duplicated enhancers. To investigate whether the shadow enhancer pair drives less noisy expression, we calculate the coefficient of variation (CV) associated with the shadow enhancer pair or either duplicated enhancer across time of nc14. **A**. The shadow enhancer pair displays lower temporal expression noise than either duplicated enhancer. Graph is mean coefficient of variation of fluorescence traces across time as a function of embryo position. **B**. The shadow enhancer pair shows the lowest expression noise, but not the highest expression levels, indicating that the lower noise is not simply a function of higher expression. Graph is average total expression during nc14 as a function of embryo position. Error bars in A and B represent 95% confidence intervals. Total number of transcriptional spots used for graphs are given in Supplementary Table 1 by construct and AP bin.

We tracked the transcriptional activity in embryos expressing MS2 under the control of the shadow enhancer pair, a duplicated proximal enhancer, or a duplicated distal enhancer (Figure 3). To measure noise associated with each enhancer, we used these traces to calculate the coefficient of variation (CV) of transcriptional activity across nc14. CV is the standard deviation divided by the mean and provides a unitless measure of noise to allow comparisons among our enhancer constructs. We then grouped these CV values by the AP position of the transcriptional spots and found the average CV at each position for each enhancer construct. All of the enhancer constructs display the lowest expression noise at the egg length of their peak expression (Figure 3A), in agreement with previous findings of an inverse relationship between mean expression and noise levels (Dar et al., 2016; Supplemental Figure 3). The shadow enhancer pair’s expression noise is almost 30% or 15% lower, respectively, than that of the duplicated proximal or distal enhancers in their positions of maximum expression.

If the primary function of shadow enhancers is only to ensure a critical threshold of expression is reached, we would not expect to also see the lower expression noise associated with the shadow enhancer pair compared to either duplicated enhancer. Furthermore, this decreased expression noise is not simply a consequence of higher expression levels, as the shadow enhancer pair produces less mRNA than the duplicated distal enhancer during nc14 (Figure 3B). The lower expression noise associated with the shadow enhancer pair suggests that it is less susceptible to fluctuations in upstream TFs than multiple identical enhancers.

### The separation of input TFs is sufficient to explain the low noise driven by the shadow enhancer pair

To explore which factors drive the difference in CVs between the duplicated and shadow enhancer constructs, we extended our model to have a single promoter controlled by two enhancers (Figure 4A). To do so, we assumed that either or both enhancers can be looped to the promoter and drive mRNA production. The rate of mRNA production when both enhancers are looped is the sum of the rates driven by the individual enhancers. We assumed that some parameters, e.g. the TF production rates and mRNA decay rate, are the same as the single enhancer case. We allowed the parameters describing the promoter-enhancer looping dynamics (the *k_on_* and *k_off_*values) to differ, depending on the enhancer’s position in the construct relative to the promoter and whether another enhancer is present. To fit the *k_on_*and *k_off_* values, we used the medians of the 10 best single enhancer parameter sets as a starting point and performed simulated annealing to refine them.

**Figure 4:**
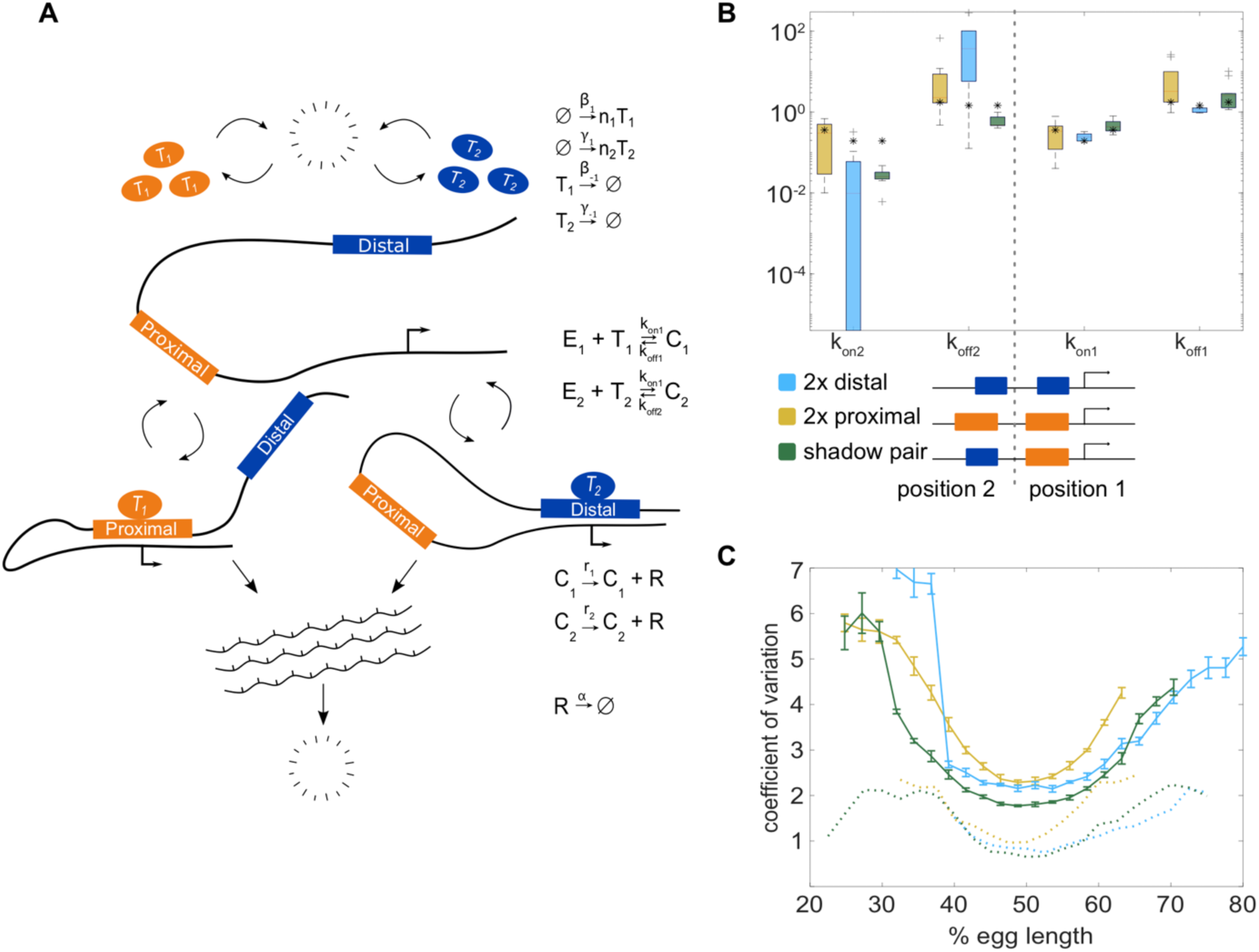
The two enhancer model recapitulates low expression noise associated with the shadow enhancer pair. To assess whether the separation of input TFs mediates the lower expression noise driven by the shadow enhancer pair, we expanded our model to incorporate two enhancers driving transcription. **A**. Schematic of the two enhancer model. We assume that when two enhancers control a single promoter, either or both can loop to the promoter and drive transcription. We defined model parameters as in Figure 2, and only allowed the *k_on_* and *k_off_*values to vary from the single enhancer model. **B**. To understand the effect of adding a second enhancer, we examined how the *k_on_* and *k_off_*values vary from those in the single enhancer model. We plotted the distribution of the values for *k_on_* and *k_off_* for each enhancer in the three different constructs measured. The distribution shows the values derived from the 10 best-fitting parameter sets, and the black star in each column indicates the *k_on_*or *k_off_* value from the corresponding single enhancer model. In general, the *k_off_* values increased relative to the single enhancer model, and the *k_on_* values decreased, indicating that the presence of a second enhancer inhibits the activity of the first. **C.** Graph of average coefficient of variation of simulated or experimental transcriptional traces as a function of egg length. The model is able to recapitulate the lower expression noise seen with the shadow enhancer pair with no additional fitting, indicating that the separation of TF inputs to the two enhancers is sufficient to explain this observation. Simulated data are shown in solid lines, experimental data are shown in dotted lines.

This approach allowed us to examine how the model parameters that describe promoter-enhancer looping dynamics change when two enhancers are controlling the same promoter. We compared the *k_off_* and *k_on_* values for each enhancer in the two enhancer constructs to their values from the single enhancer model. We generally found that *k_off_* values increased and *k_on_* values decreased (Figure 4B). The effect is most pronounced in the duplicated distal enhancer, with large changes to the *k_off_*and *k_on_* values for the enhancer in the position far from the promoter (position 2). Given that our model assumes that enhancers act additively and only allows for changes in the *k_off_* and *k_on_* values, these observed effects may indicate that either the presence of a second enhancer interferes with promoter-enhancer looping or that the promoter can be saturated. Our model cannot distinguish between these two possibilities, but these observations are consistent with our (Supplementary Figure 4) and previous results indicating that the *Kr* enhancers act sub-additively (Scholes, et al., 2019). Additionally, the dramatic changes in *k_off_* and *k_on_*values in the duplicated distal enhancer are consistent with a previous assertion that enhancer sub-additivity is most pronounced in cases of strong enhancers (Bothma et al., 2015).

We used these models to simulate transcription and predict the resulting CVs from the duplicated enhancer and shadow enhancer constructs. In line with experimental data, we found the model predicts that the shadow enhancer construct drives lower noise than the duplicated distal or duplicated proximal enhancer constructs in the middle of the embryo. This is particularly notable, as we did not explicitly fit our model to reproduce the experimentally observed CVs. There is only one fundamental difference between the shadow and duplicated enhancer models, namely the use of separate TF inputs for the shadow enhancers. Therefore, we can conclude that the separation of input TFs is sufficient to explain the low noise driven by the shadow enhancer construct.

### The shadow enhancer pair buffers against intrinsic and extrinsic sources of noise

To further validate that the more stable expression driven by the shadow enhancer pair is due to its separation of inputs, we compared the extrinsic and intrinsic noise associated with the shadow enhancer pair to that associated with either single or duplicated enhancers. To do so, we measured the transcriptional dynamics of embryos with two identical reporters in each nucleus and calculated noise sources using the approach of Elowitz, et al. (Elowitz, et al., 2002). Intrinsic noise corresponds to sources of noise, such as TF binding and unbinding, that affect each allele separately. It is quantified by the degree to which the activities of the two reporters in a single nucleus differ. Extrinsic noise corresponds to global sources of noise, such as TF levels, that affect both alleles simultaneously. It is measured by the degree to which the activities of the two reporters change together. Intrinsic and extrinsic noise are defined such that, when squared, their sum is equal to total noise^2^, which corresponds to the CV^2^ of the two identical alleles in each nucleus in our system (see Methods). Because our data do not meet one key assumption needed to measure extrinsic and intrinsic noise with the two reporter approach (see Discussion; Supplementary Figure 5), we use the terms inter-allele noise and covariance in place of intrinsic and extrinsic noise.

Based on our separation of inputs hypothesis and CV data, we expected the total noise associated with the shadow enhancer pair to be lower than that associated with the duplicated enhancers. We predicted that the shadow enhancer pair will mediate lower total expression noise through lower covariance, as the two member enhancers are regulated by different TFs. Given the complexity of predicting inter-allele noise from first principles (see Supplementary Note), we predicted that constructs with two enhancers will have lower inter-allele noise than single enhancer constructs, but did not have a strong prediction regarding the relative inter-allele noise among the different two-enhancer constructs. Comparisons of noise between the single and duplicated enhancer constructs would further allow us to discern whether reductions in noise are generally associated with two-enhancer constructs or whether this is a particular feature of the shadow enhancer pair.

Neither the duplicated proximal nor distal enhancers drive significantly lower total noise than the corresponding single enhancers, indicating that the addition of an identical enhancer is not sufficient to reduce expression noise in this system (Figure 5A). The shadow enhancer pair drives lower total expression noise than either single or duplicated enhancer, consistent with the temporal CV data in Figure 3. The median total expression noise associated with the duplicated distal and duplicated proximal enhancers is 1.4 or 2.4 times higher, respectively, than that associated with the shadow enhancer pair (Figure 5A). Note that for measurements of noise, our distal construct places the enhancer at the endogenous spacing from the promoter, as we wanted to control for positional effects on expression and noise (Scholes, et al., 2019; Supplemental Figure 6).

**Figure 5:**
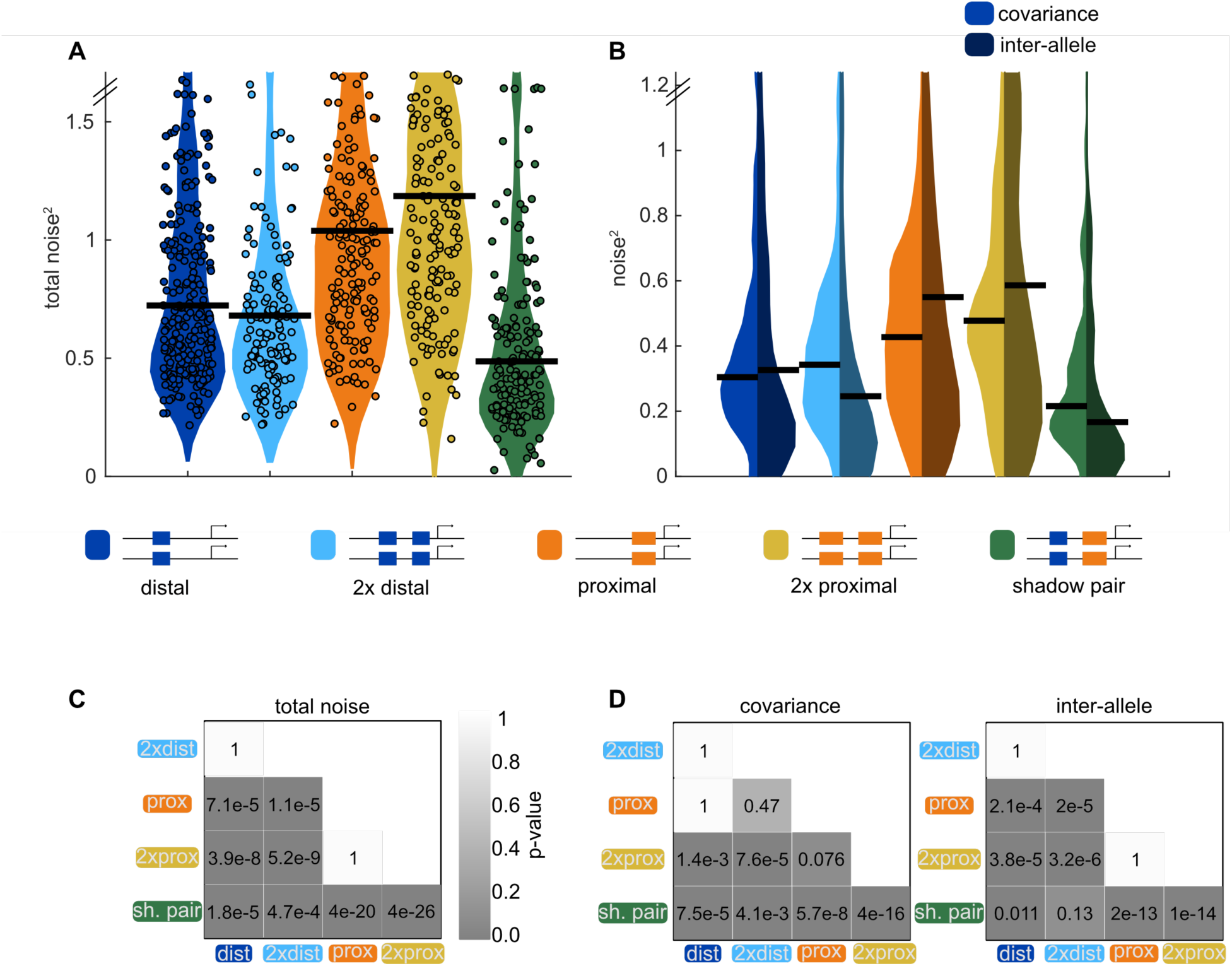
Shadow enhancer pair achieves lower total noise by buffering global and allele-specific sources of noise. To determine how the shadow enhancer pair produces lower expression noise, we calculated the total noise associated with each enhancer construct and decomposed this into the contributions of covariance and inter-allele noise. Covariance is a measure of how the activities of the two alleles in a nucleus change together and is indicative of global sources of noise. Inter-allele noise is a measure of how the activities of the two alleles differ and is indicative of allele-specific sources of noise. **A.** The shadow enhancer pair has lower total noise than single or duplicated enhancers. Circles are total noise values for individual nuclei in AP bin of peak expression for the given enhancer construct. Horizontal line represents the median. The y-axis is limited to 75th percentile of the proximal enhancer, which has the largest noise values. The shadow enhancer pair has significantly lower total noise than all other constructs. **B.** The shadow enhancer pair displays significantly lower covariance than either single or duplicated enhancer and significantly lower inter-allele noise than both single enhancers and the duplicated proximal enhancer. The left half of each violin plot shows the distribution of covariance values of nuclei in the AP bin of peak expression, while the right half shows the distribution of inter-allele noise values. Horizontal lines represent median. The y-axis is again limited to the 75th percentile of enhancer with the largest noise values, which is duplicated proximal. The lower covariance and inter-allele noise associated with the shadow enhancer pair indicates it is better able to buffer both global and allele-specific sources of noise. **C.** *p*-value table of Kruskal-Wallis pairwise comparison of the total noise values of each enhancer construct. *p*-value gradient legend applies to C and D. **D.** *p*-value table of Kruskal-Wallis pairwise comparison of covariance (on left) and inter-allele noise (on right) values for each enhancer construct. Bonferroni multiple comparison corrections were used for *p*-values in C and D. Total number of nuclei used in noise calculations are given in Supplementary Table 2.

In line with our expectations, the shadow enhancer pair has significantly lower covariance levels than either single or duplicated enhancers (Figure 5B). The shadow enhancer pair also has lower inter-allele noise than all of the other constructs, though these differences are only marginally significant (*p* = 0.13) when compared to the duplicated distal enhancer. Covariance makes a larger contribution to the total noise for the duplicated distal enhancer and the shadow enhancer pair, while inter-allele noise is the larger source of noise for the single distal enhancer and the single or duplicated proximal enhancers (Figure 5B).

The lower total noise and covariance of the shadow enhancer pair support our hypothesis that, by separating regulation of the member enhancers, the shadow enhancer pair can buffer against upstream fluctuations. The lower inter-allele noise associated with the shadow enhancer pair warrants further investigation. A simple theoretical approach predicts that two enhancer constructs will have lower inter-allele noise (see Supplementary Note). Given that this is not universally observed in our data, this suggests that there is still much to discover about how inter-allele noise changes as additional enhancers control a gene’s transcription.

### The shadow enhancer pair drives low noise at several temperatures

We showed the *Kr* shadow enhancer pair drives expression with lower total noise than either single or duplicated enhancer, yet previous studies have generally found individual member enhancers of a shadow enhancer set are dispensable under ideal conditions (Frankel, et al., 2010; Perry et al., 2011; Osterwalder, et al., 2018). However, in the face of environmental or genetic stress, the full shadow enhancer group is necessary for proper development (Frankel et al., 2010; Osterwalder, et al., 2018; Perry, et al., 2011). We therefore decided to investigate whether temperature stress causes significant increases in expression noise and whether the shadow enhancer pair or duplicated enhancers can buffer potential increases in noise.

Similar to our findings at ambient temperature (26.5°C), the shadow enhancer pair drives lower total noise than all other tested enhancer constructs at 32°C (Figure 6B). At 32°C, the duplicated distal and duplicated proximal enhancers display 35% or 52%, respectively, higher total noise than the shadow enhancer pair. At 17°C, the shadow enhancer pair has approximately 46% lower total noise than either the single or duplicated proximal enhancer, 21% lower total noise than the single distal enhancer, and is not significantly different than the duplicated distal enhancer (Figure 6A). As seen by the variety of shapes in the temperature response curves (Figure 6C), temperature perturbations have enhancer-specific effects, suggesting input TFs may differ in their response to temperature change. The low noise driven by the shadow enhancer pair across conditions is consistent with previous studies showing shadow enhancers are required for robust gene expression at elevated and lowered temperatures (Frankel, et al., 2010; Perry, et al., 2010).

**Figure 6:**
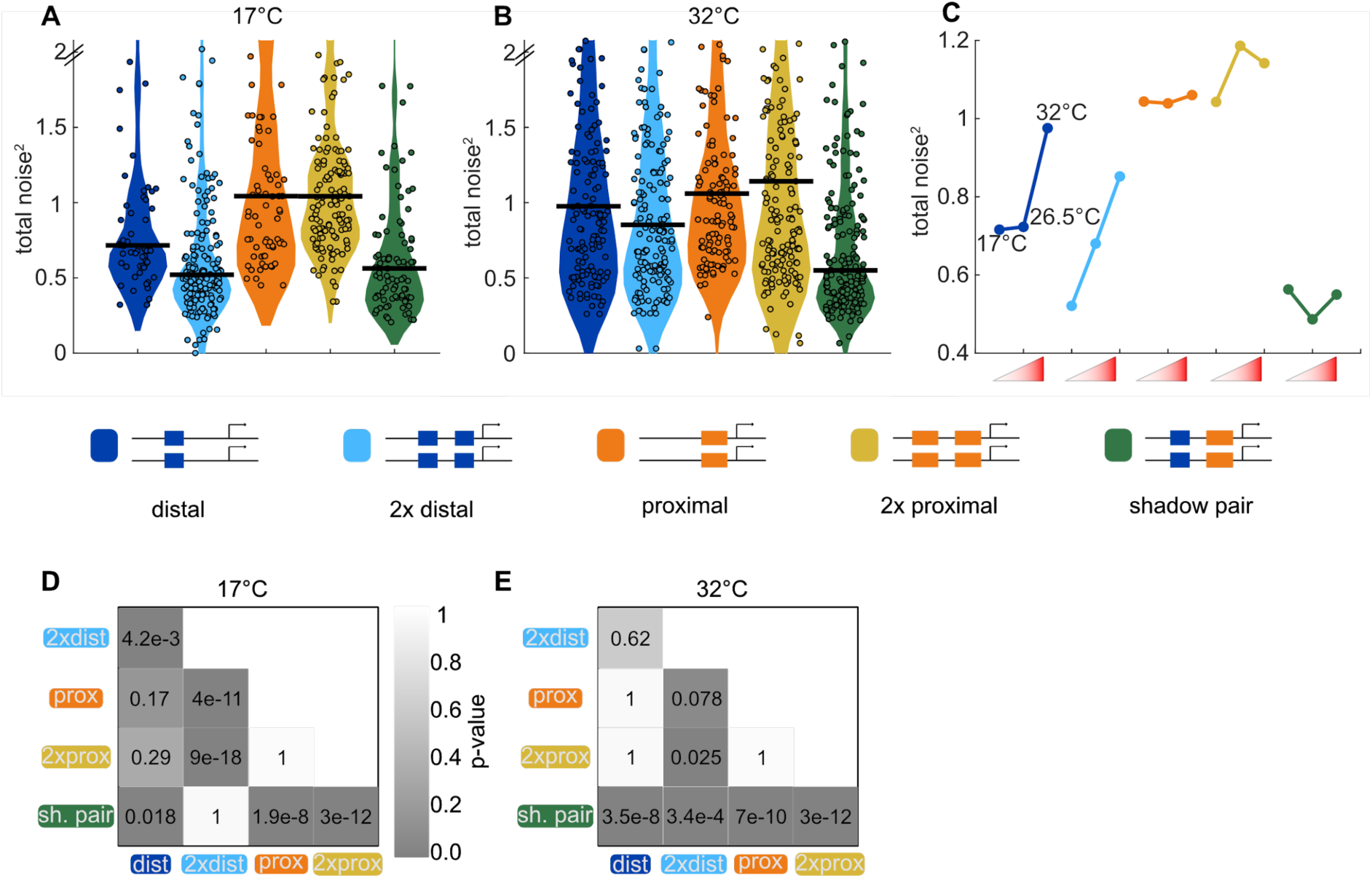
Shadow enhancer pair maintains lower total noise across temperature perturbations. To test the ability of each enhancer construct to buffer temperature perturbations, we measured the total expression noise associated with each for embryos imaged at 17°C or 32°C. **A**. The shadow enhancer pair displays significantly lower total noise than the single or duplicated proximal enhancer and the single distal enhancer at 17°C. Circles are total noise values for individual nuclei in AP bin of peak expression for the given enhancer construct and horizontal bars represent medians. The y-axis is limited to 75th percentile of construct with highest total noise at 17°C (single proximal). **B**. The shadow enhancer pair has significantly lower total noise than all other constructs at 32°C. The y-axis is limited to 75th percentile of the enhancer construct with highest total noise at 32°C (duplicated proximal). **C.** Temperature changes have different effects on the total noise associated with the different enhancers. The median total noise value at the AP bin of peak expression at the three measured temperatures is shown for each enhancer construct. Within each enhancer, the median total noise values are shown left to right for 17°C, 26.5°C, and 32°C. **D**. *p*-value table of Kruskal-Wallis pairwise comparison of the total noise values of each enhancer construct at 17°C. *p*-value gradient legend applies to D and E. **E**. *p*-value table of Kruskal-Wallis pairwise comparison of the total noise values of each enhancer construct at 32°C.

## Discussion

Fluctuations in the levels of transcripts and proteins are an unavoidable challenge to precise developmental patterning (Raser & O’Shea, 2005; Arias & Hayward, 2006; Hansen, et al., 2018). Given that shadow enhancers are common and necessary for robust gene expression (Osterwalder, et al., 2018; Frankel, et al., 2010; Perry, et al., 2010), we proposed that shadow enhancers may function to buffer the effects of fluctuations in the levels of key developmental TFs. To address this, we have, for the first time, extensively characterized the noise associated with shadow enhancers critical for patterning the early *Drosophila* embryo. By tracking biallelic transcription in living embryos, we tested the hypothesis that shadow enhancers buffer noise through a separation of TF inputs to the individual member enhancers. Our results show that TF fluctuations play a significant role in transcriptional noise and that a shadow enhancer pair is better able to buffer both extrinsic and intrinsic sources of noise than duplicated enhancers. Using a simple mathematical model, we found that fluctuations in TF levels are required to reproduce the observed correlations between reporter activity and that the low noise driven by the shadow enhancer pair is a natural consequence of the separation of TF inputs to the member enhancers. Lastly, we showed that a shadow enhancer pair is uniquely able to buffer expression noise across a wide range of temperatures. Together, these results support the hypothesis that the separation of inputs of shadow enhancers allow them to buffer input TF noise and therefore drive more robust gene expression patterns during development.

### Temporal fluctuations in transcription factor levels drive expression noise in the embryo

When measured in fixed embryos, the TFs used in *Drosophila* embryonic development show remarkably precise expression patterns, displaying errors smaller than the width of a single nucleus (Dubuis, et al., 2013; Gregor, et al., 2007; Little 2013; He, et al., 2008). It therefore was unclear whether fluctuations in these regulators play a significant role in transcriptional noise in the developing embryo. By measuring the temporal dynamics of the individual *Kr* enhancers, each of which is controlled by different transcriptional activators, we show that TF fluctuations do significantly contribute to the noise in transcriptional output of a single enhancer. Within a nucleus, expression controlled by the two different *Kr* enhancers is far less correlated than expression driven by two copies of the same enhancer, indicating that TF inputs, as opposed to more global factors, are the primary regulators of transcriptional bursting in this system.

Given that individual *Kr* enhancers are influenced by fluctuations in input TFs, it may seem puzzling that endogenous *Kr* expression patterns are rather reproducible (Little 2013). Previous work has cited the role of spatial and temporal averaging, which buffers noisy nascent transcriptional dynamics to generate more precise expression levels. Shadow enhancers operate upstream of this averaging, driving less noisy nascent transcription than either single enhancers or enhancer duplications.

### A stochastic model underscores importance of transcription factor fluctuations

We developed a stochastic mathematical model of *Kr* enhancer dynamics and mRNA production that recapitulates our main experimental results. This model is based on that by (Bothma, et al., 2015), but it is expanded to include the dynamics of a TF that regulates each enhancer. We placed a strong emphasis on the simplicity of this model, e.g. by using a single abstract TF for each enhancer. This choice both avoids a combinatorial explosion of parameters and makes the model results and parameters easier to interpret. One of the most notable features of the model is that it recreates the differences in noise between shadow and duplicated enhancer constructs without any additional fitting, indicating that these differences are a direct result of the separation of input TFs to the proximal and distal enhancers.

Future versions of this model can include refinements. For example, in the current model, we do not include the influence of repressiveTFs or fluctuations that affect transcription globally. The absence of these features may partially explain the non-zero correlation experimentally observed in the shadow heterozygote embryos. Future experiments and models can also be designed to identify the mechanism of enhancer non-additivity: changes in promoter-enhancer looping, saturation of the promoter, or other mechanisms.

### Noise source decomposition suggests competition between reporters

In our investigation of sources of noise, we decomposed total noise into extrinsic and intrinsic components as in (Elowitz, et al., 2002). In that study, the authors showed that the activity of one reporter did not inhibit expression of the other reporter, and therefore their calculations assume no negative covariance between the reporters’ expression output. In our system, we found a small amount of negative covariance between the activity of two alleles in the same nucleus (Supplemental Figure 5). For this reason, we called our measurements covariance and inter-allele noise. The negative covariance we observe indicates that activity at one allele can sometimes interfere with activity at the other allele, suggesting competition for limited amounts of a factor necessary for reporter visualization. The two possible limiting factors are MCP-GFP or an endogenous factor required for transcription. If MCP-GFP were limiting, we would expect to see the highest levels of negative covariance at the center of the embryo, where the highest number of transcripts are produced and bound by MCP-GFP. Since the fraction of nuclei with negative covariance is highest at the edges of the expression domain (Supplementary Figure 5), the limiting resource is likely not MCP-GFP, but instead a spatially-patterned endogenous factor, like a TF.

Currently, the field largely assumes that adding reporters does not appreciably affect expression of other genes. However, sequestering TFs within repetitive regions of DNA can impact gene expression (Liu, et al., 2007; Janssen, et al., 2000), and a few case studies show that reporters can affect endogenous gene expression (Laboulaye, et al., 2018; Thompson & Gasson, 2001). If TF competition is responsible for the observed negative covariance between reporters, a closer examination of the effects of transgenic reporters on the endogenous system is warranted. In addition, TF competition may be a feature, not a bug, of developmental gene expression control, as modeling has indicated that molecular competition can decrease expression noise and correlate expression of multiple targets (Yuan, et al., 2018).

### Additional functions of shadow enhancers and outlook

There are likely several features of shadow enhancers selected by evolution outside of their noise-suppression capabilities. Preger-Ben Noon, et al. recently showed that all shadow enhancers of *shavenbaby*, a developmental TF gene in *Drosophila*, drive expression patterns in tissues and times outside of their previously-characterized domains in the larval cuticle (Preger-Ben Noon, et al., 2018). This suggests that shadow enhancers, while seemingly redundant at one developmental stage, may play separate, non-redundant roles in other stages or tissues. In several other cases, both members of a shadow enhancer pair are required for the precise expression pattern generated by the endogenous locus (El-Sherif & Levine, 2016; Perry, et al., 2012; Dunipace, et al., 2011; Perry, et al., 2011; Yan, et al., 2017). In the case of *Kr*, the early embryonic enhancers drive observable levels of expression in additional tissues and time points, but these expression patterns overlap those driven by additional, generally stronger, enhancers, suggesting that the primary role of the proximal and distal enhancers is in early embryonic patterning (Hoch, et al., 1990). In addition, the endogenous expression domain of *Kr* is best recapitulated by the pair of shadow enhancers (El-Sherif & Levine, 2016). Therefore, while we cannot rule out the possibility that the proximal and distal enhancers perform separate functions at later stages, it seems that their primary function, and evolutionary substrate, is controlling *Kruppel* expression pattern and noise levels during early embryonic development.

Here, we have investigated the details of shadow enhancer function for a particular system, and we expect that some key observations may generalize to many sets of shadow enhancers. Shadow enhancers seem to be a general feature of developmental systems (Cannavo, et al., 2016; Osterwalder, et al., 2018), but the diversity among them has yet to be specifically addressed. While we worked with a pair of shadow enhancers with clearly separated TF activators, shadow enhancers can come in much larger groups and with varying degrees of TF input separation between the individual enhancers (Cannavo, et al., 2016; Osterwalder, et a., 2018). To discern how expression dynamics and noise driven by shadow enhancers depend on their degree of TF input separation, we are investigating these characteristics in additional sets of shadow enhancers with varying degrees of differential TF regulation. Our current experimental data and computational results, combined with that gathered from additional shadow enhancers will inform fuller models of how developmental systems ensure precision and robustness.

## Materials and Methods

### Generation of transgenic reporter fly lines

The single, duplicated, or shadow enhancers were each cloned into the pBphi vector, upstream of the *Kruppel* promoter, 24 MS2 repeats, and a *yellow* reporter gene as in (Fukaya, et al., 2016). We defined the proximal enhancer as chromosome 2R:25224832-25226417, the distal enhancer as chromosome 2R:25222618-25223777, and the promoter as chromosome 2R:25226611-25226951, using the *Drosophila melanogaster* dm6 release coordinates. The precise sequences for each reporter construct are given in Supplementary File 1. For the allele correlation experiments, each enhancer was cloned 192 bp upstream of the *Kr* promoter, separated by the endogenous sequence found between the proximal enhancer and the promoter. For transcriptional noise experiments, the distal enhancer was placed at its endogenous spacing, 2835 bp upstream of the promoter, and the proximal enhancer sequence was replaced by a region of the lambda genome that is predicted to have few relevant TF binding sites. In the shadow enhancer pair or duplicated enhancer constructs, the two enhancers were separated by the sequence separating the proximal and distal enhancers in the endogenous locus.

Using phiC31-mediated integration, each reporter construct was integrated into the same site on the second chromosomes by injection into yw; PBac{y[+]-attP-3B}VK00002 (BDRC stock #9723) embryos by BestGene Inc. (Chino Hills, CA). To produce embryos with biallelic expression of the MS2 reporter, female flies expressing RFP-tagged histones and GFP-tagged MCP (yw; His-RFP/Cyo; MCP-GFP/TM3.Sb) were crossed with males containing one of the enhancer-MS2 reporter constructs. Virgin female F1 offspring were then mated with males of the same parental genotype, except in the case of shadow heterozygous flies, which were mated with males containing the other single enhancer-MS2 reporter.

### Sample preparation and image acquisition

Live embryos were collected prior to nc14, dechorionated, mounted on a permeable membrane, immersed in Halocarbon 27 oil, and put under a glass coverslip as in (Garcia, et al., 2013). Individual embryos were then imaged on a Nikon A1R point scanning confocal microscope using a 60X/1.4 N.A. oil immersion objective and laser settings of 40uW for 488nm and 35uW for 561nm. To track transcription, 21 slice Z-stacks, at 0.5um steps, were taken throughout the length of nc14 at roughly 30 second intervals. To identify the Z-stack’s position in the embryo, the whole embryo was imaged after the end of nc14 at 20x using the same laser power settings. Later in the analysis, each transcriptional spot’s location is described as falling into one of 42 AP bins, with the first bin at the anterior of the embryo. Unless otherwise indicated, embryos were imaged at ambient temperature, which was on average 26.5°C. To image at other temperatures, embryos were either heated or cooled using the Bioscience Tools (Highland, CA) heating-cooling stage and accompanying water-cooling unit.

### Calculation of transcription parameters

For every spot of transcription imaged, the fluorescence traces across the time of nc14 were first subject to smoothing by the LOWESS method with a span of 10%. The resulting smoothed traces were used to measure transcriptional parameters and noise. Traces consisting of fewer than three time frames were removed from calculations. To calculate transcription parameters, we used the smoothed traces to determine if the promoter was active or inactive at each time point. A promoter was called active if the slope of its trace (change in fluorescence) between that point and the next was greater than or equal to the instantaneous fluorescence value calculated for one mRNA molecule (F_RNAP_, described below). Once called active, the promoter is considered active until the slope of the fluorescence trace becomes less than or equal to the negative instantaneous fluorescence value of one mRNA molecule, at which point it is called inactive until another active point is reached. The instantaneous fluorescence of a single mRNA was chosen as the threshold because we reasoned that an increase in fluorescence greater than or equal to that of a single transcript is indicative of an actively producing promoter, while a decrease in fluorescence greater than that associated with a single transcript indicates transcripts are primarily dissociating from, not being produced, at this locus. Visual inspection of fluorescence traces agreed well with the burst calling produced by this method (Supplemental Figure 7).

Using these traces and promoter states, we measured burst size, frequency and duration. Burst size is defined as the integrated area under the curve of each transcriptional burst. Duration is defined as the amount of time occurring between the frame a promoter is determined active and the frame it is next determined inactive. Frequency is defined as the number of bursts occurring in the period of time from the first time the promoter is called active until 50 minutes into nc14 or the movie ends, whichever is first. The time of first activity was used for frequency calculations because the different enhancer constructs showed different characteristic times to first transcriptional burst during nc14. For these, and all other measurements, we control for position of the transcription trace by first individually analyzing the trace and then using all the traces in each AP bin to calculate summary statistics of the transcriptional dynamics and noise values at that AP position.

### Conversion of integrated fluorescence to mRNA molecules

To put our results in physiologically relevant units, we calibrated our fluorescence measurements in terms of mRNA molecules. As in (Lammers, et al., 2018), for our microscope, we determined a calibration factor, α, between our MS2 signal integrated over nc13, F_MS2_,and the number of mRNAs generated by a single allele from the same reporter construct in the same time interval, N_FISH_, using the *hunchback* P2 enhancer reporter construct (Garcia et al., 2013). Using this conversion factor, we can calculate the integrated fluorescence of a single mRNA (F_1_) as well as the instantaneous fluorescence of an mRNA molecule (F_RNAP_). With our microscope, F_RNAP_ is 379 AU/RNAP and F_1_ is 1338 AU/RNAP·min. With these values, we are able to convert both integrated and instantaneous fluorescence into total mRNAs produced and number of nascent mRNAs present at a single time point, by dividing by F_1_ and F_RNAP_, respectively.

### Calculation of noise metrics

To calculate the temporal CV each transcriptional spot *i*, we used the formula:

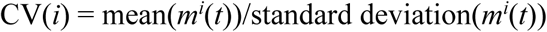

where *m^i^*(*t*) is the fluorescence of spot *i* and time *t*.

We also decomposed the total noise experienced in each nucleus to inter-allele noise and co-variance, analogous to the approach of (Elowitz, et al., 2002).

Inter-allele noise is calculated one nucleus at a time. It is the mean square difference between the fluorescence of the two alleles in a single nucleus:

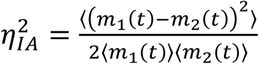

where *m_1_(t)* is the fluorescence of one allele in the nucleus at time *t*, and *m_2_(t)* is the fluorescence of the other allele in the same nucleus and the angled brackets indicate the mean across the time of nc14.

Covariance is the covariance of the activity of the two alleles in the same nucleus across the time of nc14:

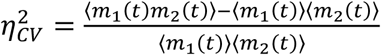

The inter-allele and covariance values are defined such that they sum to give the total transcriptional noise displayed by the two alleles in a single nucleus.

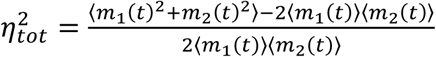

This total noise value is equal to the coefficient of variation of the expression of the two alleles in a single nucleus across the time of nc14.

### Statistical methods

To determine any significant differences in total noise, covariance, or inter-allele noise values between the different enhancer constructs, we performed Kruskal-Wallis tests with the Bonferroni multiple comparison correction.

### Description of the single enhancer model and associated parameters

We constructed a model of enhancer-driven transcription based on the following chemical reaction network,

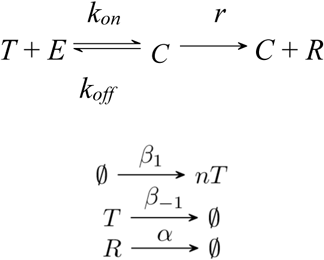

where *E* is an enhancer that interacts with a transcription factor *T*, which together bind to the promoter at a rate *k_on_* to form the active promoter-enhancer complex *C*. When the promoter is in this active form, it leads to the production of mRNA denoted by *R*, which degrades by diffusion from the gene locus at a rate *α*. Transcription is interrupted whenever the complex *C* disassociates spontaneously at a rate *k*_off_. In the bursting TFs model, the transcription factor *T* appears at a rate *β*_1_ and degrades at a rate *β*_−1_. To recapitulate *Kruppel* expression patterns, the value of *β*_1_ was assumed to be given by

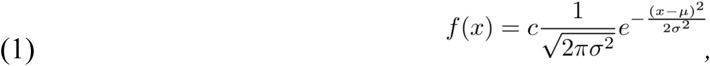

where *x* is the percentage along the length of the egg and *c* is a scaling constant. Since *Kruppel* activity peaks near the center of the egg, we chose *µ* = 50, while *c* and *σ* were fitted along with the other parameters. Lastly, *n* was assumed to be fixed across the length of the egg.

We also generated a constant TF model, which is an adaptation of the model in (Bothma et al., 2015). This model implicitly assumes that TF numbers are constant and, therefore, are incorporated into the value of *k_on_*as described by the reactions

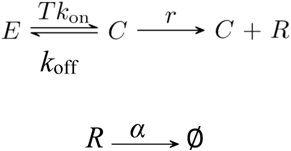

In this case, the value for *T* was fitted for each bin in a similar way to *β*_1_, i.e. the constant number of TFs was assumed to be described by equation (1) (values were rounded to the nearest integer).

To simulate the transcriptional traces, we implemented a stochastic approach. Individual chemical events such as enhancer-promoter looping take place at random times and are influenced by transcription factor numbers. Individual trajectories of chemical species over time were calculated using the Gillespie algorithm (Gillespie, 1976), and these trajectories are comparable to the experimentally measured transcriptional traces. Since the enhancer is either bound or not bound to the promoter, we imposed the constraint that *C* + *E* = 1 when simulating model dynamics.

### Estimation of model parameters from experimental data

To yield a starting estimate for the *k_on_*and *k_off_* parameters, we defined the start and end of a burst as the time when the reactions 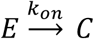 and 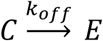 occur, respectively. The length of the *i*th burst was defined as the range of [*b_i_,p_i_*] where *b_i_* corresponds to the time of the *i*^th^ instance of the reaction 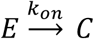 and *p_i_*to the time of the *i*^th^ instance of the reaction 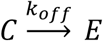. The time between the *i*^th^ burst and the *i* + 1^th^ burst is [*p_i_,b_i_*_+1_]. The Gillespie algorithm dictates that the time spent in any given state is determined by an exponentially distributed random variable with a rate parameter equal to the product of two parts: the sum of rate constants of the outgoing reactions, and the number of possible reactions. If the enhancer is either bound or unbound, we have that *C* = 1 or *E* = 1, respectively. Therefore, by letting *t_b_* be the average time between bursts and *t_d_* be the average duration of a burst, we can write

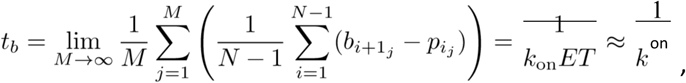

and

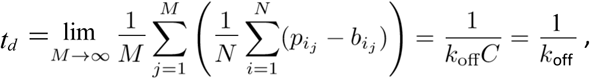

where *N* is the number of bursts for spot *j*, *b_ij_* and *p_ij_*denote the beginning and end of burst *i* in spot *j* respectively, and *M* denotes the total number of spots in the egg. The right-hand sides are given by the expected value of the exponential distribution and the assumption that, on average, T is close to 1. While this may not be the case for *T*, the assumption provides a convenient upper bound for the average time between bursts, which is likely not to have a much smaller value for a lower bound. (A low enough value of *t_b_* would imply nearly constant fluorescence intensity instead of bursts.) Finally, the average duration of a burst *t_d_* can be calculated directly from the data and used to obtain *k*_off_ by calculating 1*/t_d_*. Similarly, the average time between bursts *t_b_* is readily available from the data giving us *k*_on_ ≈ 1*/t_b_*.

We were able to directly estimate mRNA production and degradation rates from the experimental data. To estimate *α*, we focused on periods of mRNA decay; i.e. periods where no active transcription is taking place and are thus described by

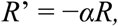

which in turn can be solved to be

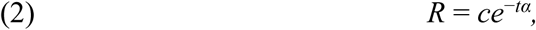

where *c* is a constant of integration. Taking the derivative of equation 2 yields

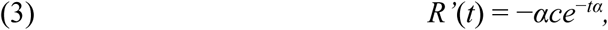

which corresponds to the slope of the decaying burst. We define the interval of decay of the *i*^th^ burst as [*p_i_,b_i_*_+1_]. For some point *t*_0_ ∈(*p_i_,b_i_*_+1_), let *R*_0_ = *R*(*t*_0_) = *ce*^−*t*0^*^α^*. Solving this expression for *c* gives that *c* = *R*_0_*e^t^*^0^*^α^*. Substituting for *c* in equation 3 evaluated at *t*_0_ results in *R*’(*t*_0_) = −*αR*_0_*e^t^*^0^*^α^e*^−*t*0^*^α^* = −*αR*_0_. Then, it follows that

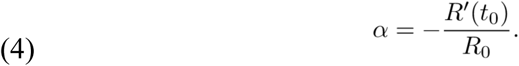

In other words, the rate of decay of mRNA fluorescence can be calculated from any trace by taking the ratio of the slope during burst decay and its intensity at a given time *t*_0_ ∈(*p_i_,b_i_*_+1_).

Adjacent measurements of fluorescence intensity from the single enhancer systems were used to approximate the slope at each point in the traces. Then, equation 4 was applied to each point. A histogram of all calculated values was generated (Supplemental Figure 8). In this figure, there was a clear peak, which provided us with an estimate of *α* ≈ 1.95.

The estimation of *r* was done for periods of active transcription, which are also accompanied by simultaneous mRNA decay. By noting that *C* = 1 during mRNA transcription, we can approximate these periods as the zeroth order process

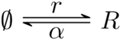

The differential equation associated with this system is given by

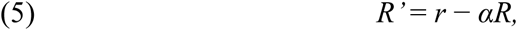

and has steady state *R*\* = *r/α*. Equation 5 can be solved explicitly for *R* by choosing

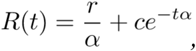

where *c* is a constant of integration. For two adjacent measurements at times *t*_1_ and *t*_2_ we can write their respective measured amounts of mRNA as

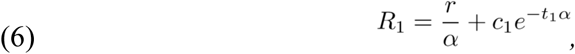

and

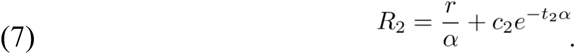

Solving for *c*_1_ and *c*_2_ gives

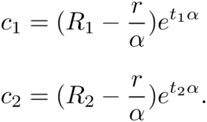

The short-term fluctuations of mRNA from *R*_1_ to *R*_2_ between two adjacent discrete time points in the stochastic system can be approximated by equations 6 and 7. This implies that

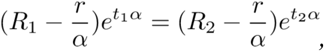

which in turn gives

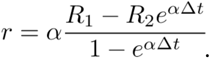

Therefore, the estimation of *r* can be computed given two adjacent measurements of fluorescence and the time between them. Finally, we used a similar approach as done with *α* to calculate values of *r* from fluorescence data. However, unlike *α*, *r* was calculated for each bin to account for differences in transcriptional efficiency across the length of the embryo.

### Parameter fitting with simulated annealing

Simulations and parameter fitting were done with MATLAB®. Optimization in fitting was done by minimizing the sum of squared errors (SSE) between the normalized vectors of burst properties and allele correlations of the experimental and simulated data. In particular, a vector *y* of experimental data was created by concatenating the following vectors: burst size, integrated fluorescence, frequency, duration, and allele correlation across the length of the embryo. The vector *y* was subsequently normalized by dividing each burst property by the largest element in their respective vectors (except correlation which by definition is unitless between -1 and 1). A vector *x* was created in an analogous fashion to *y* but using simulated instead of experimental data. However, *x* was normalized using the same elements that were used to normalize *y*. Then, the discrepancy between the experimental and simulated data was measured by

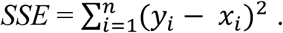

We used a high-performance computing cluster to compute 200 independent runs of parameter fitting with simulated annealing for each model variant. The algorithm requires an initial guess of the parameter set *P*_0_, an initial temperature Γ_0_, a final temperature Γ’, the number of iterations per temperature *N*, and a cooling factor *µ*. Then, each iteration is as follows:

1. If the current iteration *i* is such that *i > N*, then update the current temperature Γ*_k_* = *µ^k^*Γ_0_ to *µ^k^*^+1^Γ_0_ and set *i* = 0. Otherwise, set *i* to *i* + 1.
2. Check if Γ*_k_ <* Γ’. If so, return the current parameter set *P_j_* and terminate.
3. Choose a parameter randomly from *P_j_* and multiply it by a value sampled from a normal distribution with a mean equal to 1. The standard deviation of such distribution should be continuously updated to be Γ*_k_*. The result of this step is the newly generated parameter set *P_j_*_+1_.
4. Calculate Δ*E* as the difference in SSE between the data generated by *P_j_* and that generated by *P_j_*_+1_. Update *P_j_* to *P_j_*_+1_ if Δ*E <* 0 or with probability *p < e*^Δ*E/*Γ*k*^ where *p* is a uniformly distributed random number.
5. Repeat all steps until termination.

To generate our results, we chose Γ_0_ = 1, Γ’ = Γ_0_*/*10, *N* = 30, and *µ* = 0.8. We observed an improvement in the quality of the fittings by using analysis-derived parameter values as initial guesses instead of values given through random sampling. The sampled space ranged from 10^−3^ to 10^3^ for all parameters, except *n,* which was sampled from 10^0^ to 10^2^, and *σ,* which was randomly chosen to be an integer between 1 and 20. Equal numbers of parameter values were sampled at each order of magnitude. The analysis in the section above was used to estimate the parameters in *P*_0_. Parameters that were not estimated in the previous section were given the following initial guesses: *n* = 10, *β*_−1_ = 1, *σ* = 6, and *c* = 40. Initial guesses for *c* and *σ* were based on the experimental observation that there is little transcription outside of 20-80% egg length. Based on this observation, simulations were limited to this egg length range, as well. For the constant TFs model, both analysis-derived and random initial parameter values were used to maximize the likelihood of finding any parameter set capable of recapitulating the observed allele correlation.

### Generation of simulated experimental data

Parameter sets resulting from fitting were sorted in ascending order based on their sum of squared errors, and the 10 lowest error parameter sets are what we called the 10 best parameter sets. For all figures, we simulated 80 spots per bin and simulated each bin 5 times to generate error bars. Data for the distal enhancer at the proximal location was used to reproduce simulated allele correlations in all cases.

Gillespie simulations update the counts of each chemical species at random time intervals. However, for ease of parameter fitting and to better recapitulate the experiments, we generated data in two distinct timescales: one consisting of 30 second intervals after which mRNA counts were recorded, and another consisting of random time intervals generated by the algorithm after which chemical counts were updated. The former one was used for all parameter fitting rounds and generation of figures.

### Description of two enhancer model, parameter estimation, and fitting

To explore two enhancer systems, we expanded our previous model to include an additional enhancer. First, we considered duplicated enhancer systems, which consist of either two proximal or two distal enhancers. Enhancers were denoted by *E*_1_ and *E*_2_, which correspond to two identical enhancers that exist in different locations relative to the promoter. They are activated by the same transcription factors as described by the reactions

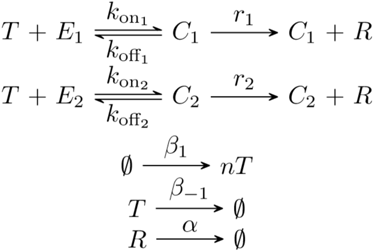

Without loss of generality, we used *E*_1_ to denote the enhancer at the proximal location and *E*_2_ to denote the enhancer at the distal location. This model describes independent enhancer dynamics; i.e. the behavior of one enhancer does not affect the behavior of the other, and, as such, both enhancers can be simultaneously looped to the promoter. Consequently, to account for potential enhancer interference or competition for the promoter, we assumed distinct *k*_on_ and *k*_off_ values for each enhancer in the duplicated enhancer constructs. We also used distinct values of *r* for each distal enhancer in the duplicated distal construct since fluorescence data was available for this enhancer at the proximal and endogenous location. For proximal enhancers, we assume *r*_1_ = *r*_2_.

To describe the dynamics of the shadow enhancer pair, we denoted the activators for *E*_1_ (the proximal enhancer) and *E*_2_ (the distal enhancer) by *T*_1_ and *T*_2_, respectively:

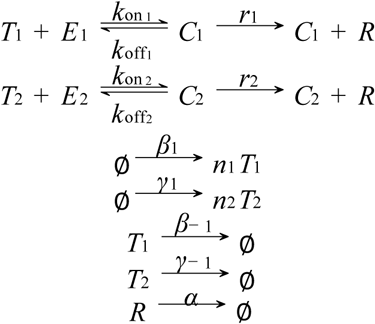

The production rate of *T_2_*, *γ*_1_, was calculated in the same way as production rate of *T_1_*, *β*_1_, but differed in the values of *c* and *σ*. The two enhancer models were also used to calculate allele correlation between homozygotes and heterozygotes because a distinction between the mRNA produced by *C*_1_ and *C*_2_ was made. This approach works because, e.g., when considering the homozygote embryos, each single enhancer resides in the same nucleus and is therefore affected by the same fluctuating TF numbers. In the duplicated enhancer model, each enhancer *E_1_*or *E_2_* is affected by the same fluctuations in the number of transcription factor T. An analogous logic applies to the heterozygotes.

To fit the two enhancer models to experimental data, we retained several parameters from the single enhancer models. Parameters *r* and *α* were directly calculated from the data, and, as such, did not vary across models. We assume that parameters concerning transcription factors (*β*_1_, *β*_−1_, *γ*_1_, *γ*_−1_, *n*_1_, and *n*_2_) are not affected by the presence of an additional enhancer. Therefore, in our model, only *k_on_*and *k_off_* are free to change. To fit the values of *k*_on1_*, k*_on2_*, k*_off1_, and *k*_off2,_ we set the other model parameters to the median values of the 10 best parameter sets in the respective single enhancer model. We then used a similar simulating annealing approach to fit the *k_on_* and *k_off_* values. We used the resulting values to simulate transcriptional traces and to calculate the predicted CV values shown in Figure 4.

## Acknowledgements

The authors would like to thank Hernan Garcia for providing advice on MS2 imaging, the MCP-GFP fly line, and a plasmid containing the MS2 reporter construct, Nick Lammers for assistance in the calibration of MS2 signal to mRNA counts, Clarissa Scholes for assistance in image processing, Lily Li for assistance in data analysis and comments on the text, and Thomas Schilling and Rahul Warrior for useful suggestions for the project.

## Supplemental Figures

**Supplemental Figure 1:**
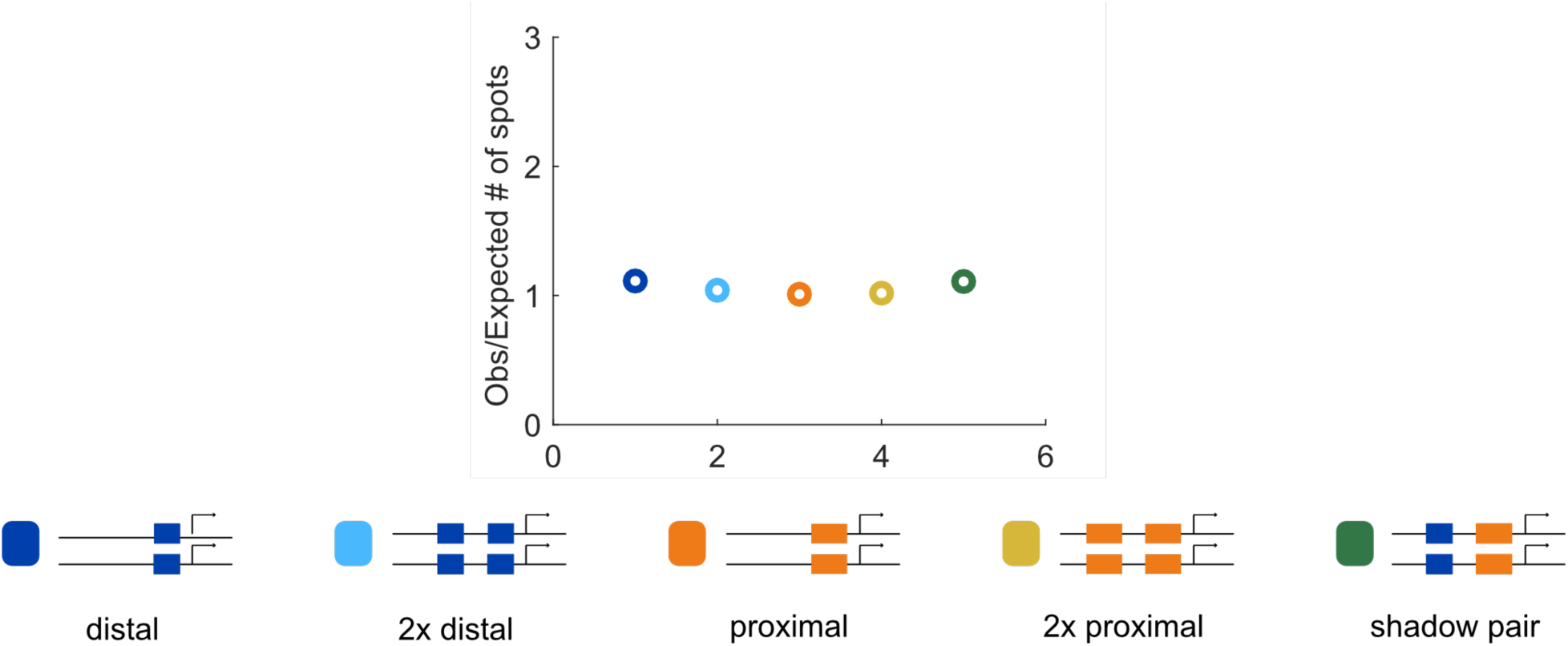
Correspondence of observed and expected number of spots. To ensure that we can accurately measure two spots of expression in the embryo, we compared the number of transcriptional spots seen in embryos hemizygous or homozygous for each construct. Our rationale was that in the absence of transvection, the number of transcriptional spots in homozygous embryos should be twice the number in embryos expressing the reporter on only one allele. The number of transcriptional spots tracked during nc14 in the AP bin of maximum expression was counted for all embryos imaged for each homozygous and hemizygous construct. The graph shows the average of this value for homozygous embryos, divided by double the value observed in the corresponding hemizygous construct. Assuming no transvection occurs, this value should be close to 1. The ratio of observed to expected number of spots is close to 1 for all of our enhancer constructs, indicating we are reliably able to track the two individual spots of transcription in single nuclei.

**Supplemental Figure 2:**
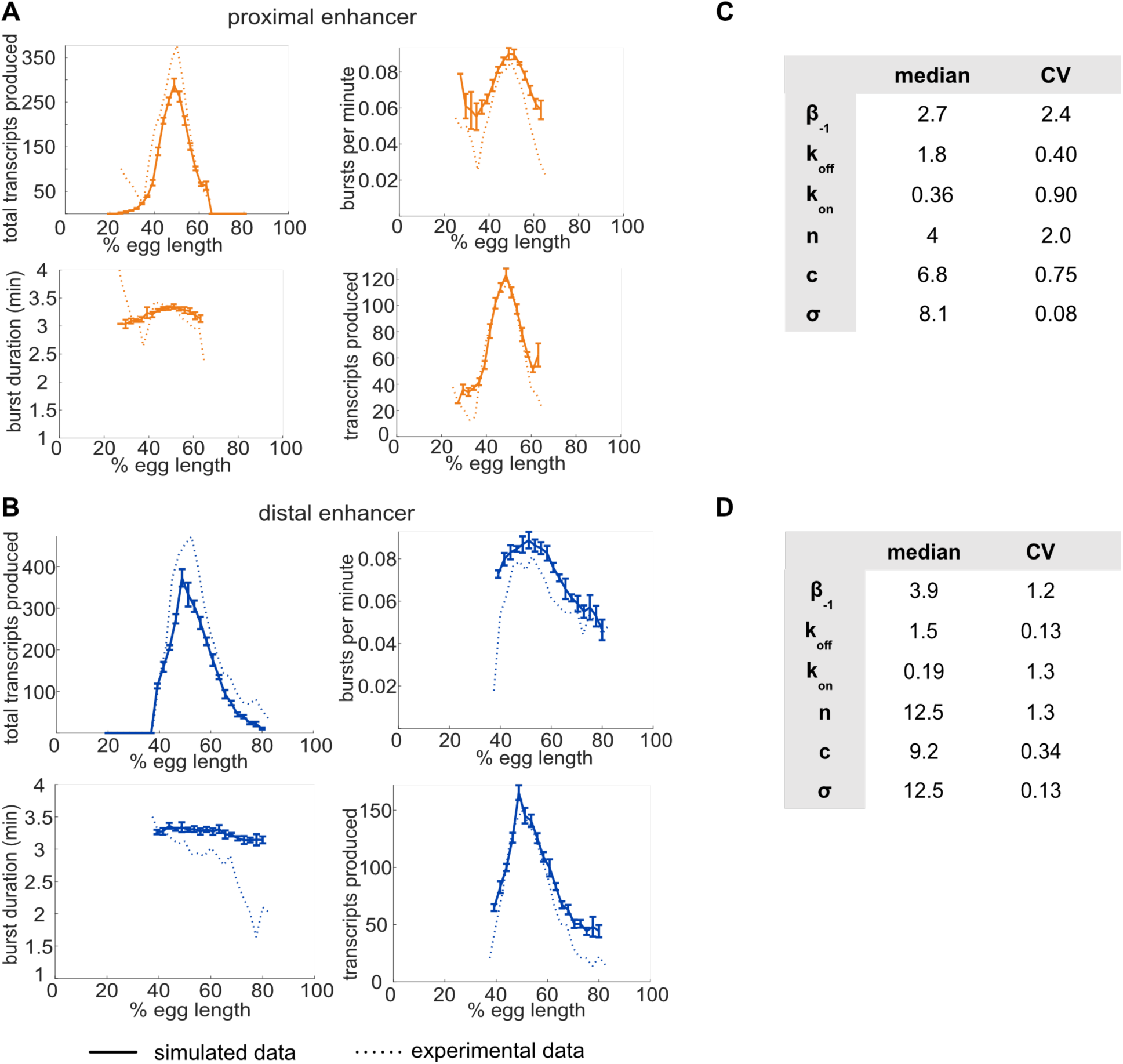
Single enhancer models recreate observed transcriptional bursting properties. To investigate whether our model is accurately simulating our experimental system, we compared the transcriptional burst properties produced by model simulations of transcription to those observed experimentally (see Supplementary Figure 7 for description of burst properties). **A**. Graphs of average values of transcriptional burst properties, total mRNA produced during nc14, burst frequency, burst duration, and burst size associated with the proximal enhancer as a function of egg length. In A and B, simulated data are represented with solid lines and experimental data are shown with dotted lines. **B**. Graphs of average values of transcriptional burst properties as in A, associated with the distal enhancer. For both the proximal and distal enhancers, our model is largely able to recapitulate the experimentally observed transcriptional burst properties associated with each enhancer. **C**. The median and CV values of the model parameters for the proximal enhancer in the top 10 performing parameter sets. **D**. The median and CV values of the model parameters for the distal enhancer in the top 10 performing parameter sets. Explanations of model parameters are given in the Methods.

**Supplemental Figure 3:**
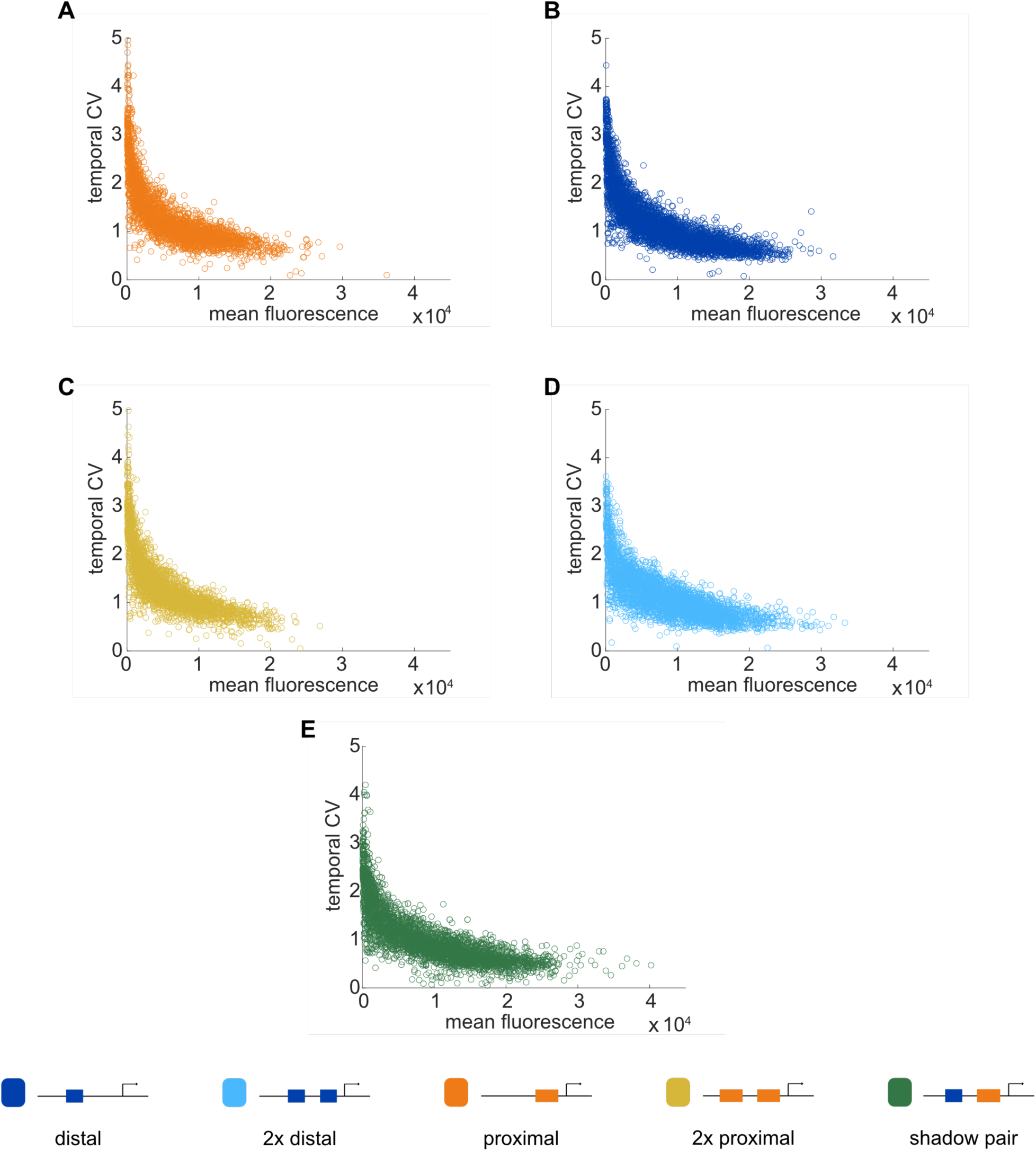
Temporal CV as a function of mean fluorescence. To investigate the relationship between our noise measurement of temporal CV and the mean activity of each construct, we plotted the temporal CV of each transcription spot as a function of its mean fluorescence. **A**. Distal; **B**. Proximal; **C**. 2x Proximal; **D**. 2x Distal; **E**. Shadow pair. With all constructs, we find the general trend that CV decreases with increasing average expression, flattening out at a baseline noise level specific to each enhancer construct.

**Supplemental Figure 4:**
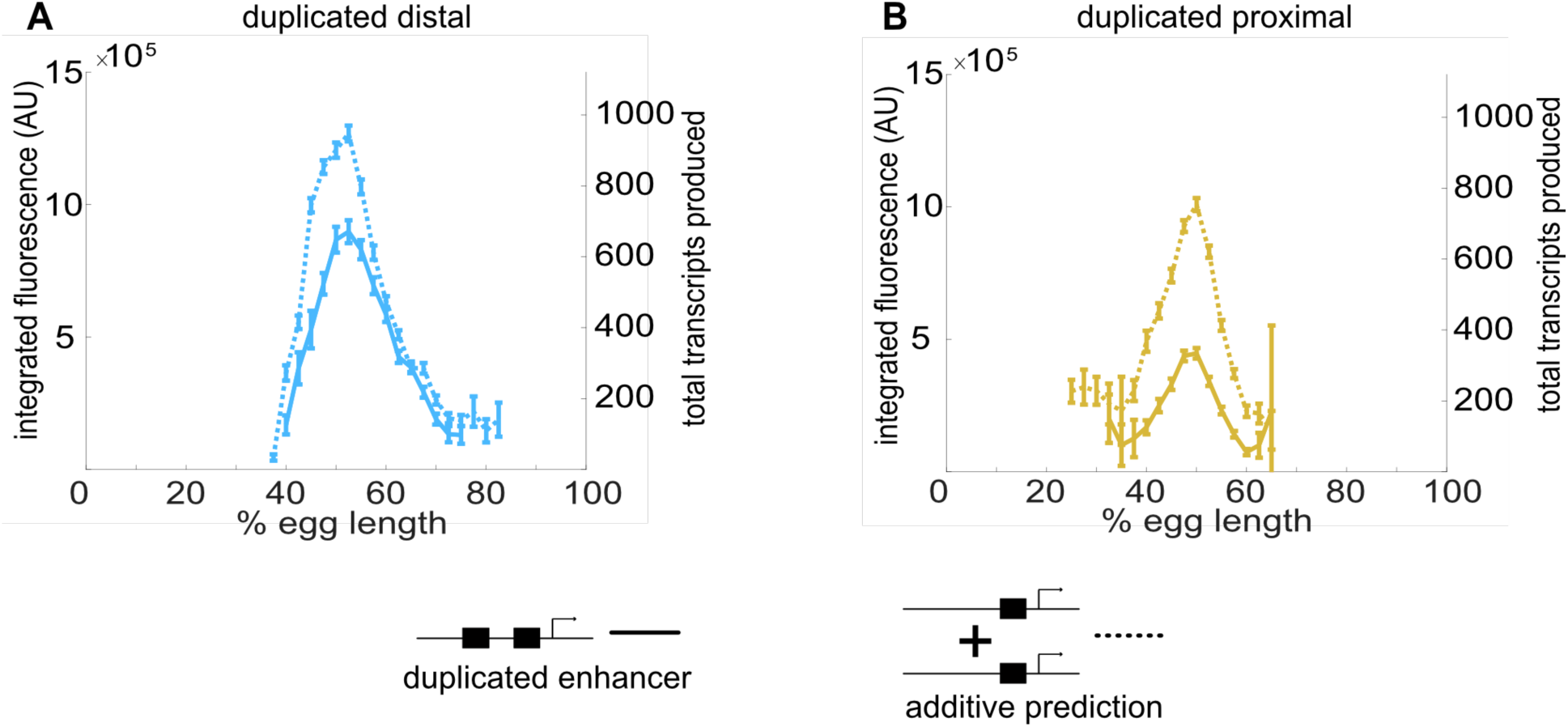
Individual *Kr* enhancers display sub-additive behavior. To assess the way input from two enhancers is integrated at the Kr promoter, we compared the experimentally observed mRNA production of duplicated enhancers to that predicted from additive behavior of the single enhancers. **A**. The duplicated distal enhancer displays sub-additive behavior. The solid line is the experimentally observed total mRNA produced by the duplicated distal enhancer during nc14 as a function of egg length and the dotted line is that expected by doubling the total mRNA produced by the single distal enhancer. **B**. The duplicated proximal enhancer also acts sub-additively. The solid line is the experimentally observed total mRNA produced by the proximal enhancer during nc14 as a function of egg length and the dotted line is that expected by doubling the total mRNA produced by the single proximal enhancer. These results, along with the observation that *k_off_* values increased and *k_on_* values decreased in our model with the addition of a second enhancer, suggests that the *Kr* enhancers compete with each other for interactions with the promoter.

**Supplemental Figure 5:**
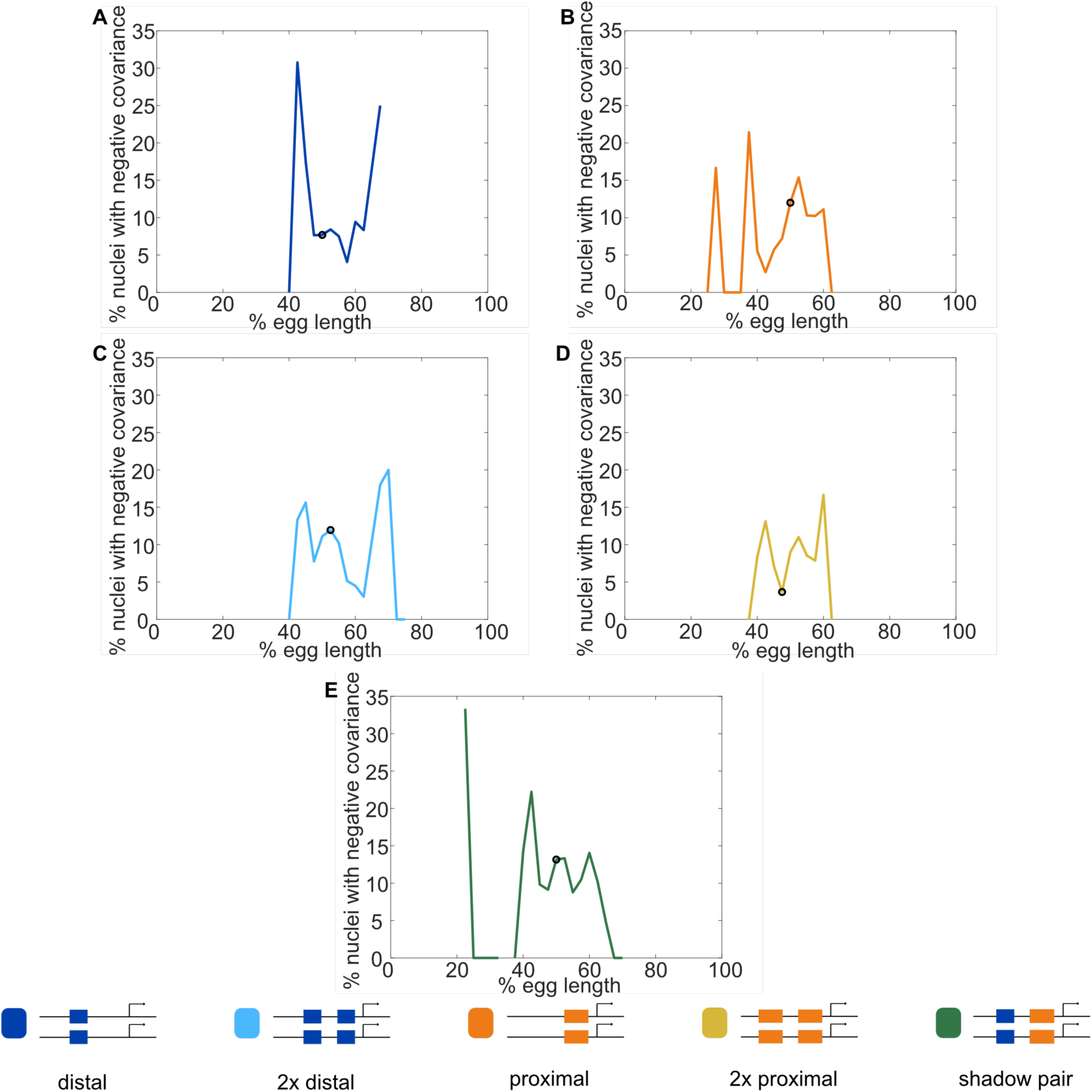
Fraction of nuclei with negative covariance of allele activity. To identify the likely cause of the observed negative covariance between allele activity in some nuclei, we calculated the fraction of nuclei displaying negative covariance out of all nuclei that had active reporter transcription. Graphs show the fraction of transcribing nuclei with negative covariance as a function of egg length for each reporter construct, with a black circle indicating the position along the embryo of maximal expression for that construct. **A**. Distal; **B**. Proximal; **C**. 2x Proximal; **D**. 2x Distal; **E**. Shadow pair. Note that for all constructs, the highest rates of negative covariance are outside of the region of maximal reporter expression. MCP-GFP is expressed uniformly along the length of the embryo and we would therefore expect if MCP-GFP were the limiting factor, we would see the highest rates of negative covariance in the center of the expression pattern, where the highest number of transcripts are produced. Instead, the highest rates of negative covariance are seen at the edges of the *Kr* expression pattern, suggesting a spatially patterned factor, such as a TF, may be what is limiting.

**Supplemental Figure 6:**
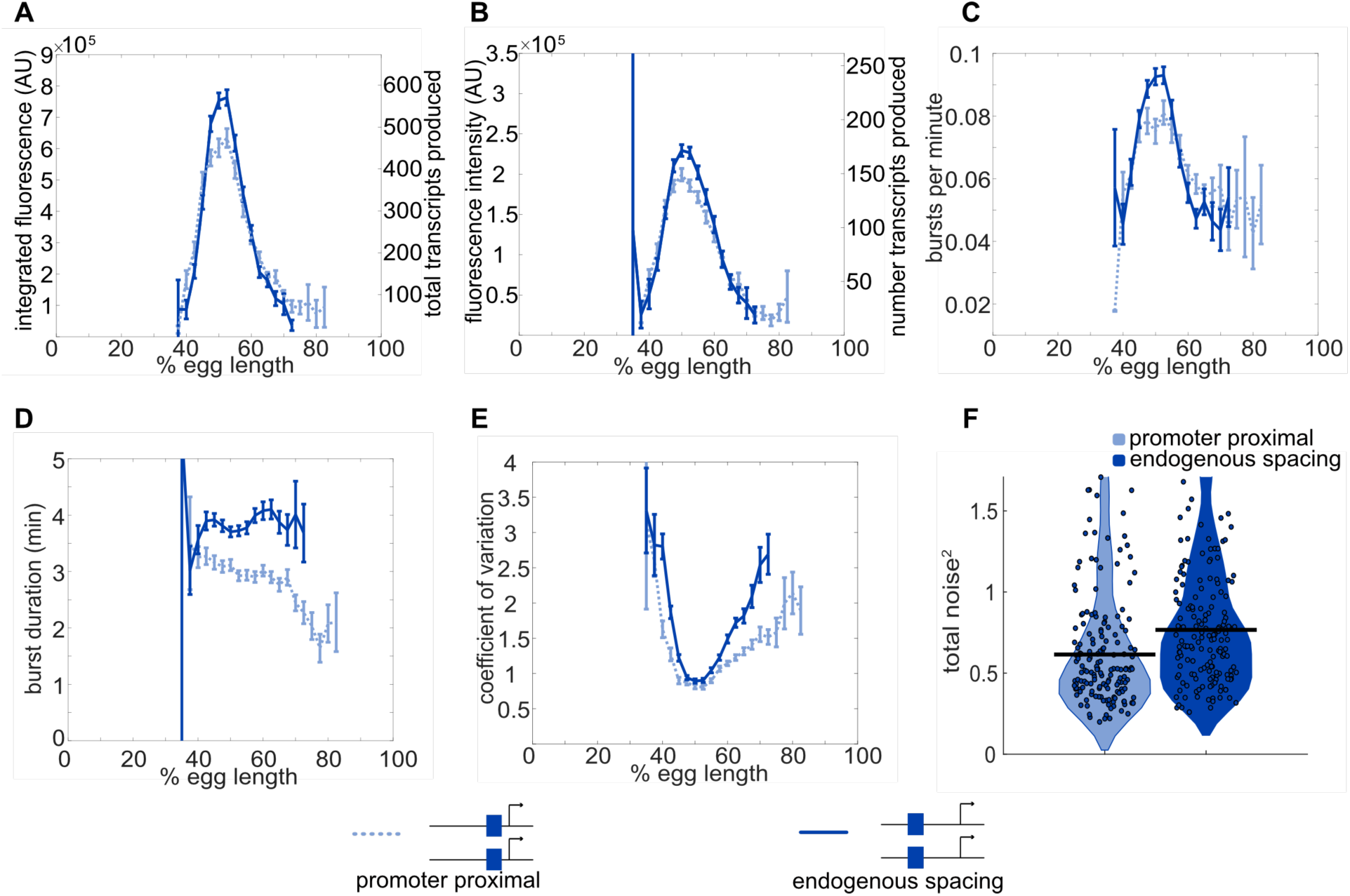
Position-dependent effects on distal enhancer. To best mimic the endogenous system, we looked at expression driven by the distal enhancer at its endogenous spacing from the promoter for our noise calculations. In this construct we replaced the sequence of the proximal enhancer with sequence of the same length from the lambda phage genome predicted to have low number of *Drosophila* TF binding sites. This increased distance from the promoter had observable effects on the transcriptional dynamics and noise associated with the distal enhancer. **A**. Comparison of total transcriptional expression mediated by the distal enhancer at its endogenous spacing or proximal to the promoter. The distal enhancer at its endogenous spacing, shown as the solid line, produces significantly more total mRNA in the center region of expression than the distal enhancer proximal to the promoter, shown as the dotted line. **B**. Comparison of the average number of transcripts produced per transcriptional burst by each distal enhancer configuration as a function of egg length. **C**. Average burst frequency associated with either distal enhancer configuration as a function of egg length. **D**. Average burst duration associated with either distal enhancer configuration as a function of egg length. **E**. Coefficient of variation of transcriptional activity across nc14 for each distal enhancer configuration as a function of egg length. **F**. Total expression noise associated with either distal enhancer configuration at the AP bin of that construct’s peak expression. The total noise distribution for the distal enhancer proximal to the promoter is on the left and that for the distal enhancer at its endogenous spacing from the promoter is on the right. The distal enhancer at its endogenous spacing displays significantly higher total noise (*p* = 0.018) than the distal enhancer proximal to the promoter. Each circle represents the total noise of an individual nucleus and the horizontal bar marks the median total noise value. Y-axis limited to the 75th percentile of the construct with the highest total noise values (distal promoter at endogenous spacing).

**Supplemental Figure 7:**
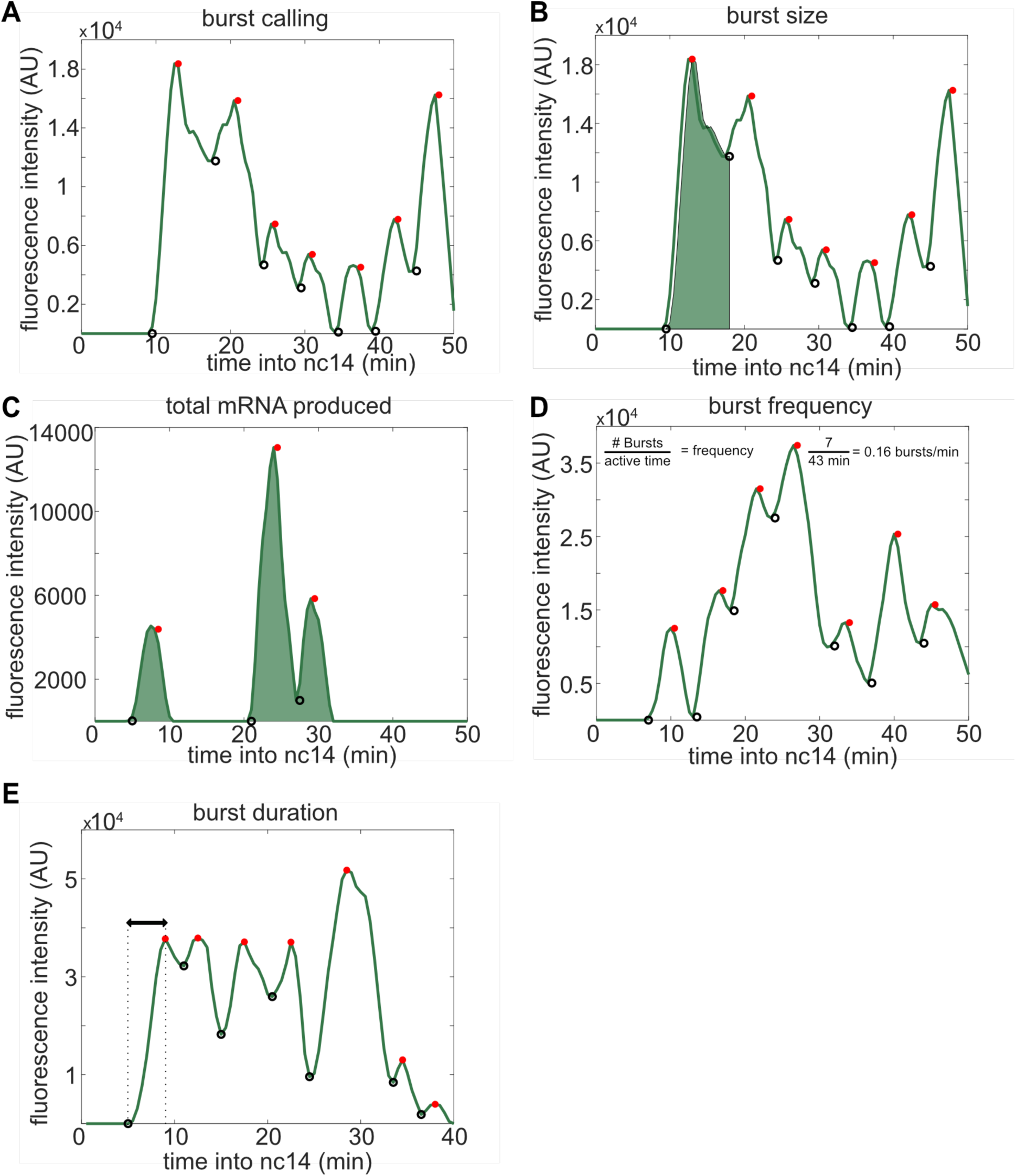
Visual inspection of burst calling algorithm. To extract the bursting parameters examined (burst size, frequency, and duration), individual fluorescence traces were first smoothed using the LOWESS method with a span of 0.1. Our burst calling algorithm then determined the periods of promoter activity or inactivity based on the slope of the fluorescence trace. **A**. Representative fluorescence trace of a single spot across the time of nc14. Black open circles indicate time points where the promoter is called “on”, red filled circles indicate time points where the promoter is called “off”. **B**. Same trace as in A with shading representing the area under the curve used to calculate the size of the first burst. This area is calculated using the trapz function in MATLAB and is done for each burst, from the time point the promoter is called “on” until the next time it is called “on”. C-E show additional representative fluorescence traces of single transcriptional spots across the time of nc14. **C**. A trace with shading representing the area under the entire curve during nc14 used to calculate the total amount of mRNA produced. This area is calculated using the trapz function in MATLAB and is done from the time the promoter is first called active until 50 minutes into nc14 or the movie ends, whichever comes first. **D**. Burst frequency is calculated by dividing the number of bursts that occur from the time the promoter is first called active until 50 minutes into nc14 or the movie ends, whichever comes first. **E**. Burst duration is defined as the amount of time between when the promoter is called active and it is next called inactive.

**Supplemental Figure 8:**
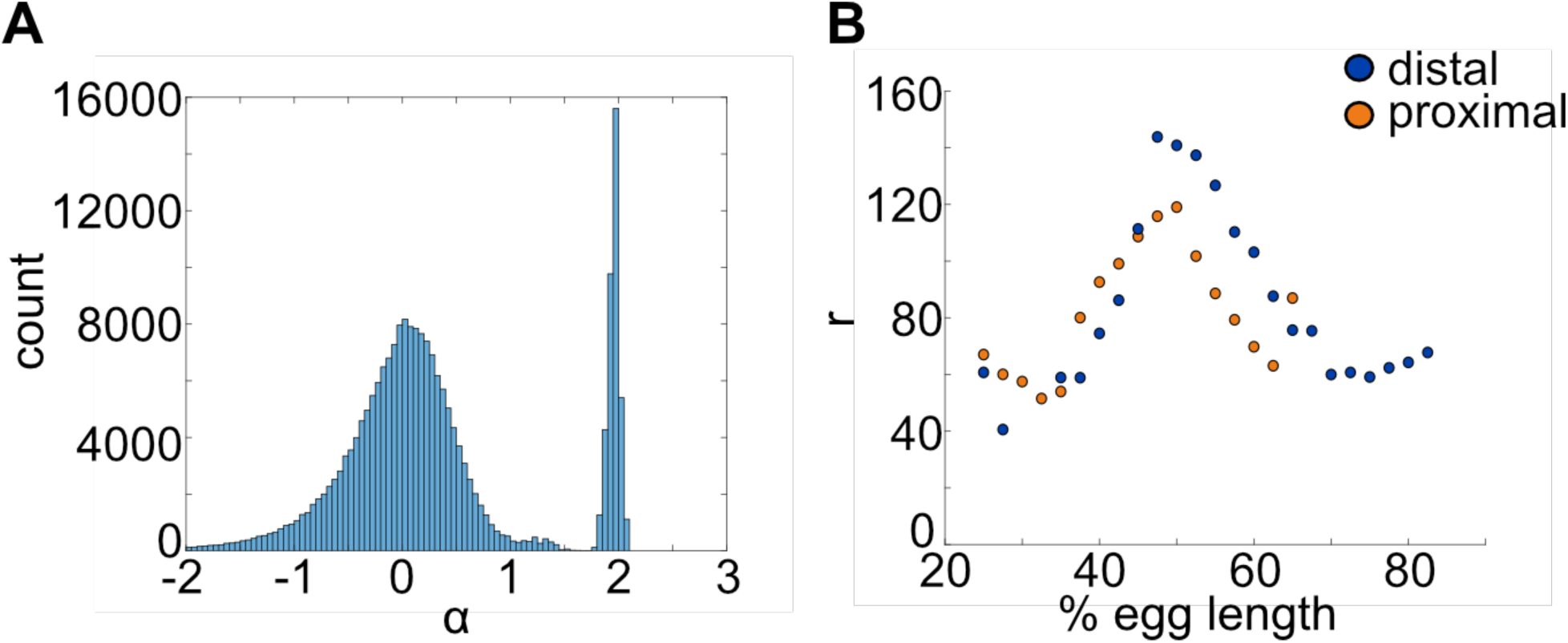
mRNA production and decay rates can be directly estimated from experimental data. The mRNA degradation parameter α and production parameter *r* were measured directly from fluorescence data without any input from the model. **A**. To estimate α, we used adjacent measurements of fluorescence intensity to approximate the slope at each point in the fluorescence traces. These values are compared with an exponential rate of mRNA decay (see Methods) and the resulting predicted values are shown in the histogram. Periods of mRNA production have negative α values and periods of decay have positive values. The histogram shows a distinct peak for α > 0, which provided us with an estimate of α ≈ 1.95. **B.** A similar computational approach was used to calculate values of *r* from fluorescence data (see Methods). We calculated different values of *r* for each bin to account for differences in transcriptional efficiency across the length of the embryo due to factors that are not explicitly included in the model. For example, different combinations of TF bound to the enhancer may give rise to different mRNA production rates. Different values of *r* were found for the proximal and distal enhancers. Notice that distal *r* values shown correspond to the distal enhancer at the proximal location.

## Additional Supplementary Materials

**Supplementary Note:** A note describing the theoretical estimates of inter-allele noise in single and two enhancer constructs.

**Supplementary Table 1:**
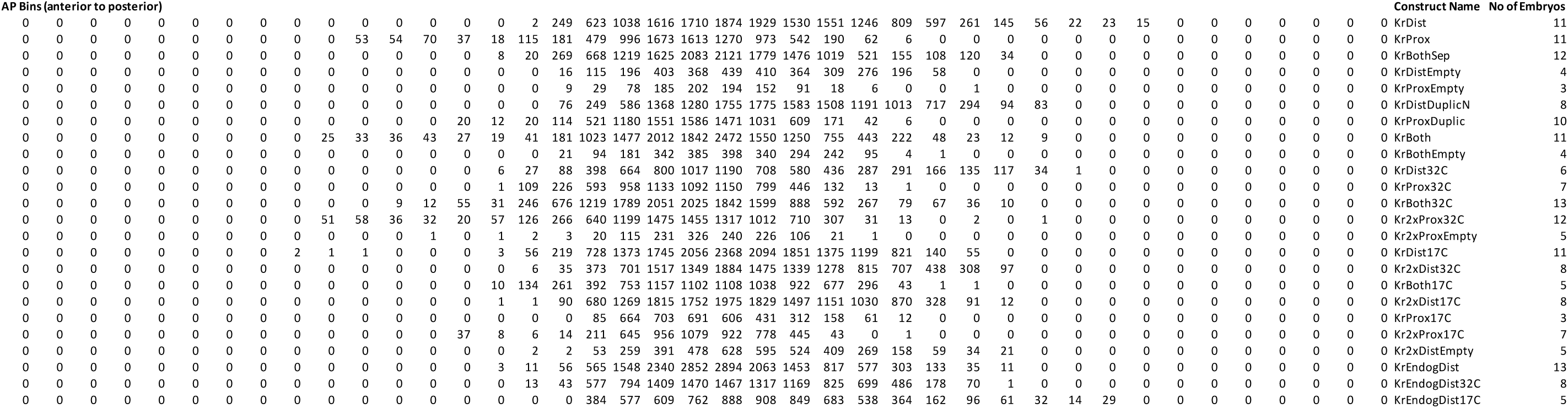
Number of total single alleles tracked for each construct. Each row corresponds to a construct, named in column 42, and columns 1-41 correspond to that AP bin of the embryo. The value in each cell in columns 1-41 is the number of single transcriptional spots used in calculations of burst size, frequency, and duration and CV in that AP bin for the given construct. The value in column 43 is the total number of independently imaged embryos for that construct.

**Supplementary Table 2:**
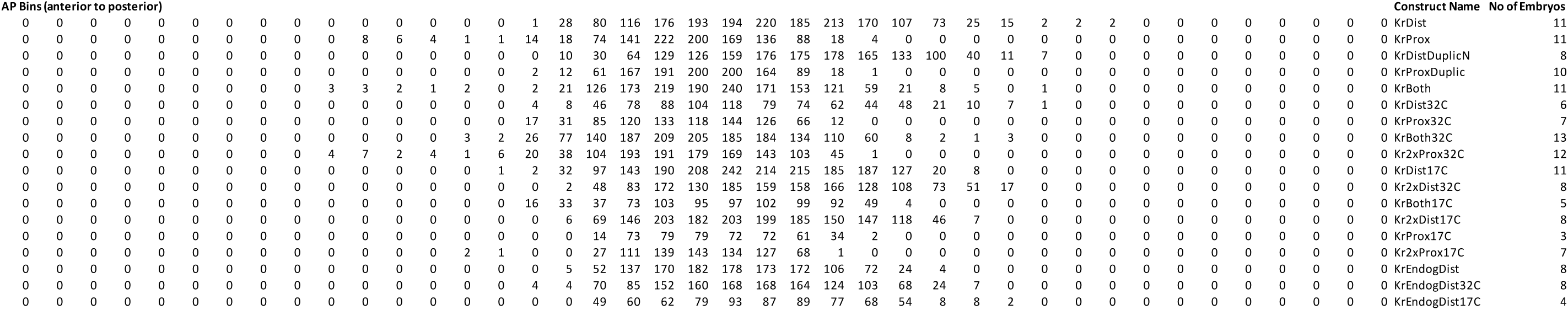
Number of nuclei tracked for each construct. Each row corresponds to a construct, named in column 42, and columns 1-41 correspond to that AP bin of the embryo. The value in each cell in columns 1-41 is the number of nuclei used for correlation and total noise/covariance/inter-allele noise calculations in that AP bin for the given construct. The value in column 43 is the total number of independently imaged embryos for that construct.

**Supplementary File 1:** The sequences of all the enhancer constructs generated in this paper.

### Supplementary Note

To make a prediction about the expected change in inter-allele noise between single and two enhancer reporter constructs, we used the theory put forth in (Sánchez and Kondev, 2008; Sanchez et al., 2011). This formalism can be used to calculate the expected mean and variance of the transcriptional output of a promoter, given the possible states of the promoter, transition rates between states, and the rate of transcription resulting from each state. In these papers, the authors apply their formalism to different promoter architectures. Here, we generate a simpler model, in which we abstract away the individual transcription factor (TF) binding configurations, which would be numerous and poorly parametrized, and simply define states by whether an enhancer is looped to the promoter and activating transcription. Since these models do not account for fluctuations that would contribute to extrinsic noise, e.g. fluctuations in TF or RNA polymerase levels, they can predict the dependence of intrinsic noise on enhancer arrangement.

To apply this model to our system, we use theses parameters:

|  |  |
| --- | --- |
| $\gamma$ | degradation rate of mRNA |
| $p$ | production rate of mRNA |
| $k$ | on rate for enhancer-promoter looping |
| $l$ | off rate for enhancer-promoter looping |

Below, we describe several models that represent different configurations of either one or two enhancers controlling a single promoter and provide the variables, as defined in (Sanchez et al., 2011), needed to calculate the coefficient of *intrinsic* variation (CV) associated with each model. Briefly, ***R*** and ***r*** describe the production rates of mRNA in the different promoter-enhancer staets, and ***K*** contains the transition rates in and out of states. Key assumptions are that the parameters describing this system are independent of both the position of the enhancer relative to the promoter and the presence of a second enhancer controlling the same promoter. We chose to make these simplifying assumptions to give the reader a general sense of the expected behavior of noise when adding an additional enhancer, since the possible behaviors are nearly infinite with the removal of these simplifying assumptions.

#### Model 1: Single enhancer

In this model, there is a single enhancer controlling one promoter.

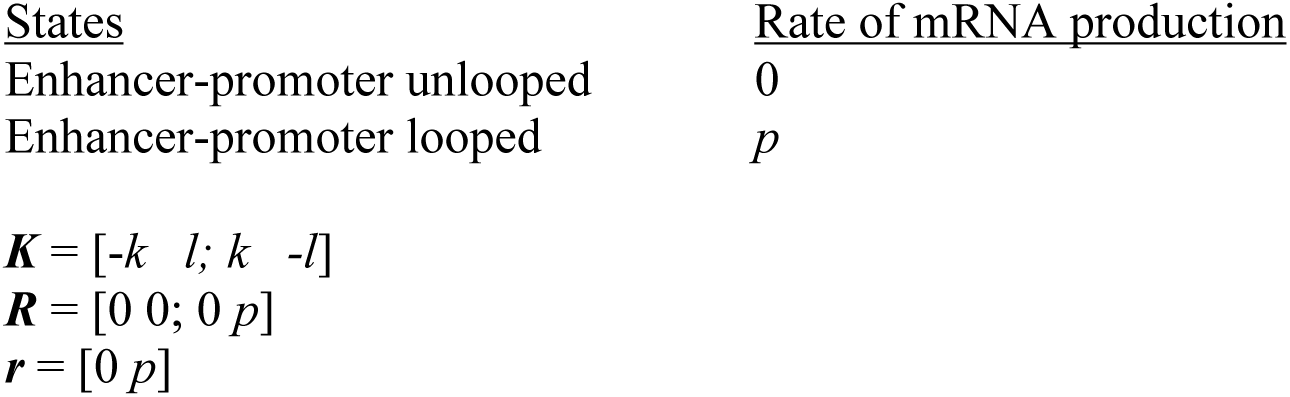

#### Model 2: OR model

In this model, there are two enhancers controlling one promoter, transcription is activated if either enhancer is looped, and both enhancers can’t be bound at the same time.

| <u>States</u> | <u>Rate of mRNA production</u> |
| --- | --- |
| Enhancers-promoter unlooped | 0 |
| Enhancer 1-promoter looped | $p$ |
| Enhancer 2-promoter looped | $p$ |

#### Model 3: Additive model

In this model, there are two enhancers controlling one promoter, transcription is activated if either enhancer is looped, and, if both enhancers are bound, transcription occurs at twice the rate of single enhancer looping states.

| <u>States</u> | <u>Rate of mRNA production</u> |
| --- | --- |
| Enhancers-promoter unlooped | 0 |
| Enhancer 1-promoter looped | $p$ |
| Enhancer 2-promoter looped | $p$ |
| Both enhancers looped | $2p$ |

#### Model 4: Synergistic model

In this model, there are two enhancers controlling one promoter, transcription is activated if either enhancer is looped, and, if both enhancers are bound, transcription occurs at three times the rate of single enhancer looping states.

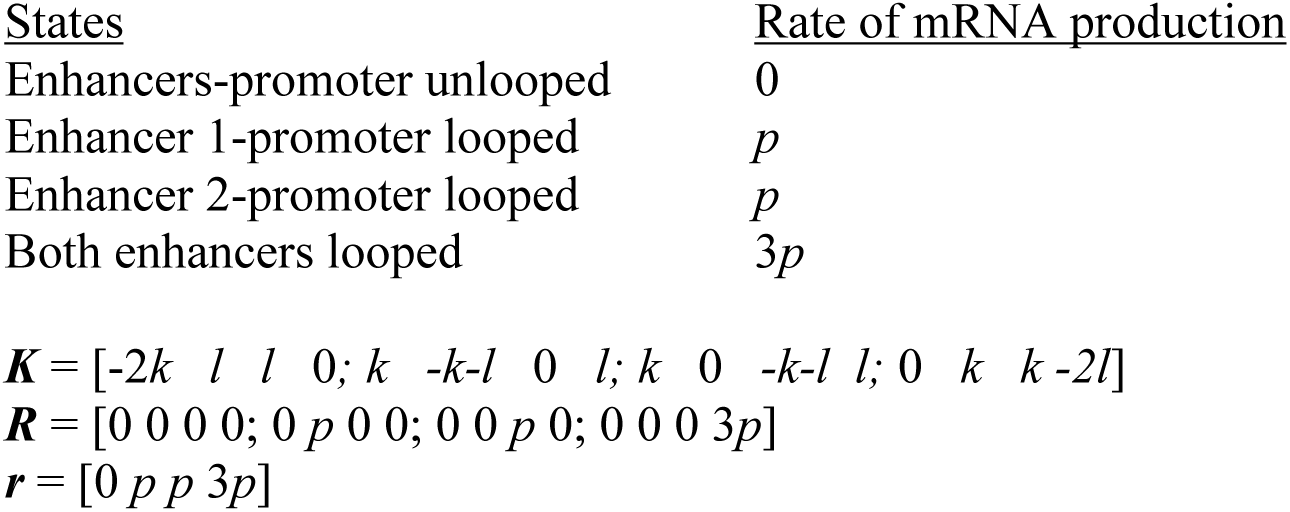

| <u>States</u> | <u>Rate of mRNA production</u> |
| --- | --- |
| Enhancers-promoter unlooped | 0 |
| Enhancer 1-promoter looped | $p$ |
| Enhancer 2-promoter looped | $p$ |
| Both enhancers looped | $3p$ |

#### Model 5: XOR model

In this model, there are two enhancers controlling one promoter, transcription is activated if either enhancer is looped, and, if both enhancers are bound, no transcription occurs.

| <u>States</u> | <u>Rate of mRNA production</u> |
| --- | --- |
| Enhancers-promoter unlooped | 0 |
| Enhancer 1-promoter looped | $p$ |
| Enhancer 2-promoter looped | $p$ |
| Both enhancers looped | 0 |

To explore the behavior of CV in these different models, we use several approaches.

**Figure 1:**
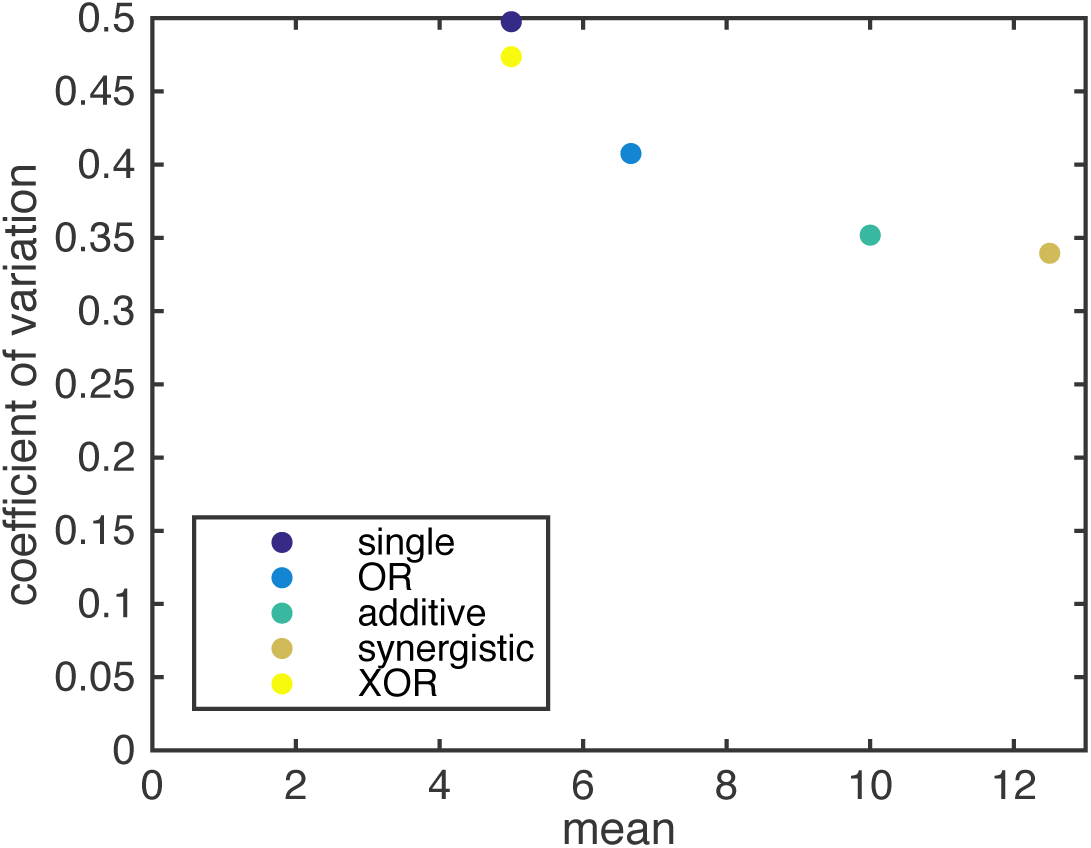
CV decreases upon the addition of a second enhancer. Here we plot the mean expression level versus CV for the five models above and one set of parameters, *k* = *l* = 1, *p* = 1, γ = 0.1. The single enhancer model (dark purple) drives the highest CV, indicating that, under the assumptions of our models, adding an additional enhancer generally lowers intrinsic noise. Except for XOR model (yellow), all other models produce more mRNA than the single enhancer model. The other colors are: blue, OR model; green, additive model; brown, synergistic model.

**Figure 2:**
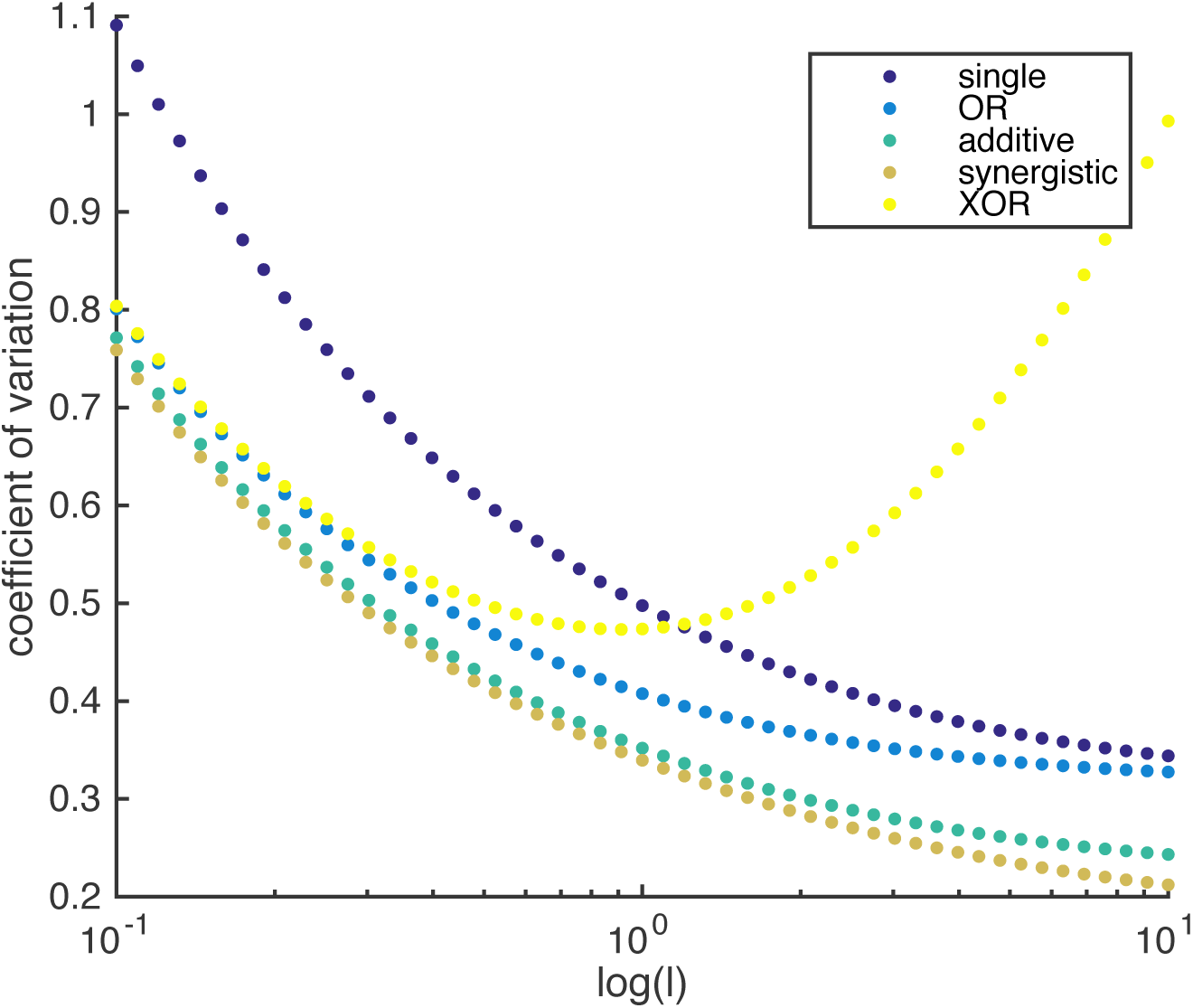
In most cases, two enhancer models drive lower noise than the single enhancer model. Here we plot the CV as a function of *l*, the rate of promoter-enhancer dissociation, for the five models above and vary *l* from 0.1 to 10 on a logarithmic scale with *k* = 1, *p* = 1, γ = 0.1. With the exception of the XOR model with low *l*, the single enhancer model drives a higher CV than the models with two enhancers for the same value of *l*.

The results above show that, under the simplifying assumptions that the production rates and on-off rates of enhancers are independent of the position and number of enhancers, the addition of a second enhancer generally lowers the predicted intrinsic noise. In our experimental data (Figure 5), we only observe a significant decrease in interallele noise for the shadow enhancer pair compared to the single distal or single proximal enhancer. Duplications of either the proximal or distal enhancer do not have significantly lower noise than their respective single enhancer constructs. Therefore, we expect that the simple addition of an identical enhancer likely does not fulfill the simplifying parameter assumptions used here and suggests that further investigation is needed to understand the complexity of the relationship between interallele noise and the numbers of enhancers controlling a promoter.

